# The L1CAM SAX-7 is an antagonistic modulator of Erk Signaling

**DOI:** 10.1101/2024.09.14.613091

**Authors:** Melinda Moseley-Alldredge, Caroline Aragón, Marcus Vargus, Divya Alley, Nirali Somia, Lihsia Chen

**Affiliations:** Department of Genetics, Cell Biology & Development, University of Minnesota, Minneapolis, MN 55455; Developmental Biology Center, University of Minnesota, Minneapolis, MN 55455

**Keywords:** *sax-7* L1CAM, MAPK, KSR-1, RanBPM, vulval development, *C. elegans*

## Abstract

L1CAMs are immunoglobulin superfamily cell adhesion molecules that ensure proper nervous system development and function. In addition to being associated with the autism and schizophrenia spectrum disorders, mutations in the L1CAM family of genes also underlie distinct developmental syndromes with neurological conditions, such as intellectual disability, spastic paraplegia, hypotonia and congenital hydrocephalus. Studies in both vertebrate and invertebrate model organisms have established conserved neurodevelopmental roles for L1CAMs; these include axon guidance, dendrite morphogenesis, synaptogenesis, and maintenance of neural architecture, among others. In *Caenorhabditis elegans*, L1CAMs, encoded by the *sax-7* gene, are required for coordinated locomotion. We previously uncovered a genetic interaction between *sax-7* and components of synaptic vesicle cycle, revealing a non-developmental role for *sax-7* in regulating synaptic activity. More recently, we determined that *sax-7* also genetically interacts with extracellular signal-related kinase (ERK) signaling in controlling coordinated locomotion. *C. elegans* ERK, encoded by the *mpk-1* gene, is a serine/threonine protein kinase belonging to the mitogen-activated protein kinase (MAPK) family that governs multiple aspects of animal development and cellular homeostasis. Here, we show this genetic interaction between *sax-7* and *mpk-1* occurs not only in cholinergic neurons for coordinated locomotion, but also extends outside the nervous system, revealing novel roles for SAX-7/L1CAM in non-neuronal processes, including vulval development. Our genetic findings in both the nervous system and developing vulva are consistent with SAX-7/L1CAM acting as an antagonistic modulator of ERK signaling.

## INTRODUCTION

The L1 family of cell adhesion molecules (L1CAMs) is a set of immunoglobulin transmembrane proteins essential in the development and function of the nervous system. Studies using vertebrate and invertebrate model organisms have uncovered conserved neurodevelopment roles for L1CAMs that include neurite outgrowth, axon guidance and myelination, dendrite morphogenesis, neuronal migration, synaptogenesis, and maintenance of neural organization (Chen and Zhou 2010; Sakurai 2012; Hortsch *et al*. 2014; Sundararajan *et al*. 2019; Gronska-Peski *et al*. 2020; Duncan *et al*. 2021). In agreement with these neuronal roles, variants in three of the four genes encoding L1CAMs (*L1CAM*, *NRCAM*, *NFSC*, and *CHL1*) result in neurodevelopmental disorders. For example, variants in the *L1CAM* gene result in the X-linked L1 Syndrome, the symptoms of which include intellectual disability, spastic paraplegia, and congenital hydrocephalus (Jouet *et al*. 1994; Vits *et al*. 1994; Van Camp *et al*. 1996; Vos and Hofstra 2010). More recently, variants in *NRCAM* and *NFASC* were identified as underlying neurodevelopmental syndromes that have both common and distinct manifestations as the L1 Syndrome. Patients with bi-allelic variants in *NRCAM* present with developmental delay, hypotonia, neuropathy, and hydrocephalus while patients with bi-allelic variants in *NFASC* present with ataxia, neuropathy, and intellectual disability (Smigiel *et al*. 2018; Monfrini *et al*. 2019; Kurolap *et al*. 2022; Elahi *et al*. 2023).

In addition to the syndromic disorders, L1CAMs are also linked to several complex genetic conditions, including Hirschsprung’s disease, the Autism and Schizophrenia spectrum disorders, addiction susceptibility, and diabetes mellitus (Parisi *et al*. 2002; Sakurai *et al*. 2002; Tam *et al*. 2010; Wallace *et al*. 2010; Ayalew *et al*. 2012; Taneera *et al*. 2012; Shaltout *et al*. 2013; Zhong *et al*. 2015; Wang and Camilleri 2019; Gauntner *et al*. 2021). Genes linked to polygenic disorders typically confer small effects; thus uncovering functional effects of each genetic association is generally quite challenging. One approach to identifying the contribution of a particular gene to a polygenic disorder is via gene modifier studies. Indeed, it was through genetic modifier studies in mice that uncovered a role for L1CAM in facilitating ganglion precursor cell migration and differentiation, thus bringing insight into how pathogenic variants in the *L1CAM* gene might contribute to Hirschsprung’s Disease (Anderson *et al*. 2006; Turner *et al*. 2009). Studies in mice also revealed mutations in *Nrcam* as genetic modifiers in murine models of peripheral neuropathy. Homozygous loss-of-function *Nrcam* mice show no overt neuromuscular functional deficits despite a delay in the formation of nodes of Ranvier in peripheral axons. However, *Nrcam* null mutations synergize with mutations in the *Lpin1*, *Sh3tc2* and *Gars* genes to cause severe neuromuscular dysfunction in the respective double mutant mice that present as a spectrum ranging from tremors to progressive paralysis; these phenotypes are modest or absent in single mutant mice (Douglas *et al*. 2009; morelli *et al*. 2017).

*Caenorhabditis elegans*, a premiere model organism to conduct genetic modifier screens, possesses a single canonical L1CAM-encoding gene, *sax-7* (Chen *et al*. 2001; Chen and Zhou 2010). SAX-7 shares many conserved neurodevelopment roles as mammalian L1CAMs (Wang *et al*. 2005; Dong *et al*. 2013; DÍaz-Balzac *et al*. 2015; Sundararajan *et al*. 2019; Cebul *et al*. 2020; Desse *et al*. 2021). Through genetic modifier screens, we previously uncovered a novel non-developmental role for *sax-7* in modulating synaptic activity. Here, *sax-7* interacts with genes that function in the synaptic vesicle cycle, such as *rab-3* and *unc-13,* to cause synthetic or enhanced locomotory abnormalities (Opperman *et al*. 2015). Reducing ERK (extracellular signal-related kinase) activity suppresses these abnormalities, revealing a role for the *mpk-1* gene, which encodes *C. elegans* ERK, in *sax-7-*mediated locomotion (Moseley-Alldredge *et al*. 2022).

ERK is a serine/threonine protein kinase belonging to the family of MAPKs (Mitogen-Activated Protein Kinase) that typically functions downstream of receptor tyrosine kinases activated by extracellular growth factors. Here, the activated receptor tyrosine kinase triggers, via the Ras GTPase, the sequential activation of a module of three kinases, RAF, MEK, and ERK (Thomas and Huganir 2004; Miningou and Blackwell 2020). ERK signaling controls multiple cellular processes, including cell proliferation and animal development. In the nervous system, ERK functions to regulate synaptic activity and plasticity (Sweatt 2004; Mao and Wang 2016; Ojea Ramos *et al*. 2022). In the *C. elegans* nervous system, ERK signaling has been shown to influence chemotactic and foraging behavior, as well as locomotory behaviors controlled by heterotrimeric Gq protein signaling (Chen *et al*. 2011; Tomida *et al*. 2012; Hamakawa *et al*. 2015; Coleman *et al*. 2018).

In this study, we examined the genetic interplay between ERK signaling and *sax-7* in the regulation of coordinated locomotion in *C. elegans*. We provide genetic evidence that points to elevated ERK signaling as contributing to the locomotion abnormalities exhibited by *sax-7* null animals. We also identified an unanticipated link between elevated ERK signaling and SAX-7 outside the nervous system, uncovering novel non-neuronal roles for SAX-7, including in vulval development. Neuronal and vulval phenotypes reflective of elevated MPK/ERK signaling in *sax-7* null backgrounds point to SAX-7 as an antagonistic modulator of MPK-1/ERK signaling.

## MATERIALS AND METHODS

### Strains

*C. elegans strains*, provided by the *Caenorhabditis* Genetics Center, were grown on nematode growth medium (NGM) plates at 21°C. N2 Bristol served as the wild-type strain (Brenner 1974). The alleles used in this study are listed by linkage groups as follows:

LG I: *arTi85 (De La Cova et al. 2017)*; *gid-1(tm3703)* (Consortium 2012)

LG II: *let-23(sy1)* (Aroian and Sternberg 1991); *narSi2* (Robinson-Thiewes *et al*. 2021)

LG III: *mpk-1(ga117)* (Lackner and Kim 1998)

LG IV: *sax-7(eq1)* (Wang *et al*. 2005); *sax-7(eq22); sax-7(eq23); sax-7(eq28)* (Moseley-Alldredge *et al*. 2022); *lin-45(sy96)* (Sternberg *et al*. 1993); *let-60(n1046)* (Beitel *et al*. 1990)

LG X: *ksr-1(ok786)* (Consortium 2012)

### The strains generated in this study are as follows

LH1195: *eqIs5*

LH1196*: sax-7(eq1); eqIs5*

LH 1445: *mpk-1(ga117)/qC1; sax-7(eq1)*

LH1446*: mpk-1(ga117)/qC1; eqIs5*

LH1447*: mpk-1(ga117)/qC1; sax-7(eq1); eqIs5*

LH1128: *sax-7(eq22) let-60(n1046)*

LH1249: *sax-7(eq1) let-60(n1046)*

LH1318: *sax-7(eq28) let-60(n1046)*

LH1140: *sax-7(eq1) let-60(n1046); ksr-1(ok786)*

LH1314: *let-23(sy1)* (*him-5(e1490)* was crossed out from the PS21 strain)

LH1182: *let-23(sy1); sax-7(eq22)*

LH1400: *let-23(sy1); sax-7(eq1)*

LH1448: *let-23(sy1); sax-7(eq1); eqEx621[Plin-31::sax-7L]*

LH1449: *sax-7(eq1); eqEx621[Plin-31::sax-7L]*

LH1450*: let-23(sy1); sax-7(eq1); eqEx622[Plin-29::sax-7L]*

LH1404: *arTi85; let-23(sy1)*

LH1410: *arTi85; let-23(sy1); sax-7(eq1)*

LH1411: *arTi85; sax-7(eq1)*

LH1443: *arTi85; let-23(sy1); sax-7(eq28)*

LH1362: *sax-7(eq22) lin-45(sy96)*

LH1434: *narSi2; mpk-1(ga117); sax-7(eq1)*

LH1451: *narSi2; sax-7(eq1)*

LH1306: *lin-31(n301); sax-7(eq1)*

LH1307: *lin-31(n301); sax-7(eq28)*

LH1089: *gid-1(tm7303)* outcrossed 3X

LH1359: *gid-1(tm7303); let-23(sy1)*

LH1423: *gid-1(tm7303); let-23(sy1); sax-7(eq1)*

LH1444: *gid-1(tm3703); sax-7(eq1)*

### Transgenes generated

*eqEx621:* pLC788 *(P_lin-31_::sax-7L,* 50ng/ul), *P_str-1_::gfp (1.5 ng/ul)*

*eqEx622:* pLC791 *(P_lin-29_::sax-7L,* 50ng/ul), *P_str-1_::gfp (1.5 ng/ul)*

*eqIs5*: pBC31 (50 ng/µl, gift of M. Ailion) was injected to generate transgenic animals carrying extrachromosomal arrays; this multicopy array was then integrated into the genome using a CRISPR/Cas9 technique (Moseley-Alldredge *et al*. 2022).

### Plasmids generated

pLC788: For VPC-directed expression of SAX-7L (*P_lin-31_::sax-7L*).

pLC791: For AC-directed expression of SAX-7L (*P_lin-29_::sax-7L*).

Both pLC788 and pLC 791 were generated by Gibson assembly of PCR-generated promoter fragments along with GFP operon sequence inserted in to a SAX-7L expression vector (pLC227) (Zhou *et al*. 2008) using the following promoter-specific primer sequences:

P*_lin-31_*F: CCAGGGCTCCTACTGGGCGG

P*_lin-31_*R: TTCAGGGAATATGTATAGAGTTTTG

P*_lin-29_*F: CGGTAGGTATGGAGAGTTG

P*_lin29_*R: ATTGCGTTGAAGAAGTTG.

pCL771: prey construct with the GID-1 cDNA subcloned into pGADT7 vector between the NdeI and XhoI sites by ProFacgen (www.profacgen.com).

### Locomotion Assay

*Crawling and radial locomotion assays* were performed using young adult animals transferred onto 60 mm NGM plates (n~30). After two minutes of recovery time post transfer, the animals were recorded at 7.5 frames per second (fps) for one minute using the WormLab Imaging System (MBF Bioscience, VT, USA). Both the x,y coordinates of individual worms and the amplitudes of their waveform were generated using WormLab. The mean amplitude represents the average displacement of the centroid along the y-axis within one cycle. Positional data was exported into the Chemotaxis and Migration Tool v2.0 (Ibidi GmbH, Martinsried, Germany), in which radial graphs were plotted. Directness of the worm trajectory is calculated by dividing the straight-line distance from the starting point by the total distance travelled.

*Swimming assays* were performed using young adult hermaphrodites transferred into a depression slide containing 1 ml M9 buffer as described (MILLER *et al*. 1996a; Opperman *et al*. 2015). After one minute of recovery time, the animals were video recorded for one minute. The number of times an animal thrashed (bending at mid body) was counted manually by examining the video recording.

### Quantifying Clr animals

10 animals per plate were examined over a span of 5 days, starting at the embryo stage. Animals that assume a Clr and transparent appearance as viewed under the dissecting microscope were transferred onto another plate to verify the Clr phenotype and to determine their outcome (eg. immobile or death) in the ensuing days.

### Quantifying excretory pore protrusions

L4 staged larvae were examined under Differential Interference Contrast Microscopy for the presence of a protuberance at the excretory pore. Depending how much the protrusion is, the phenotype is qualitatively classified as “mild” (a slight protrusion), “moderate” (a strong protrusion), and “severe” (robust protrusion with apparent tissue/debris spilling out).

### Vulval Development Analysis

The analyses were based on previously established protocols described in Gauthier and Rocheleau (2017). Briefly, L4-stage larvae semi-synchronized by time after egg-laying were mounted on 2% agarose pads, anesthetized using 10mM levamisole in M9 buffer and examined using DIC microscopy or fluorescence microscopy with an Axioplan 2 microscope (Carl Zeiss). Images are acquired using an AxioCam MRM and AxioVision 4.5 software (Carl Zeiss).

#### Analysis via vulva morphology

Mid L4-stage larvae with a complete vulval induction exhibit a single invagination along the midline that have a characteristic “Christmas tree” morphology. L4-stage larvae are identified as lacking vulval induction if their ventral midline does not have an invagination and if all VPC daughters are present. L4-stage larvae with partial vulval induction are pinpointed as those with a smaller ventral midline invagination and more than the typical number of uninduced VPC daughter cells. L4 stage larvae with elevated vulval induction are distinguished as animals with a single invagination along the midline that have a characteristic “Christmas tree” morphology as well as adjacent smaller invaginations.

#### VPC induction score

The vulvae of L4-stage larvae in *let-23* genetic backgrounds carrying the *arTi85* transgene (De La Cova *et al*. 2017) were examined via both DIC and fluorescence microscopy. The *arTi85* expresses mCherry::HIS-11 in VPCs and their descendents, providing the ability to identify which VPC is induced. In VPCs where one of the two daughters is induced is given a score of 0.5; if both daughters are induced, the VPC is given a score of 1. Hence, complete vulval induction as observed in wild-type animals is given an induction score of 3.

### SAX-7 expression

L3-staged *sax-7(eq23)* animals were mounted on 2% agarose pads and anesthetized using 10mM levamisole in M9 buffer. Images were acquired on a Nikon Ti2 inverted confocal microscope using NIS elements (Nikon Inc., Melville, NY). The image shown on Fig 3Bii is a sum slice projection obtained using ImageJ.

### GAL4-based Yeast-two-hybrid Assay

DNA constructs used:

pLC209: bait construct with the coding sequence for SAX-7 cytoplasmic tail subcloned into pGBKT7 (Zhou *et al*. 2008).

pCL771: prey construct with the GID-1 coding sequence was subcloned into pGADT7 vector between the NdeI and XhoI sites; construct generated by ProFacgen (www.profacgen.com).

#### Assay

The Yeast-two-hybrid protein interaction assay was performed by ProFacgen, using standard protocol. Briefly, Y2H Gold yeast strain was transformed with both bait and prey constructs and cultured on SD minimal media lacking specified amino acids (−L/T: leucine and tryptophan or −L/T/H/A: leucine, tryptophan, histidine, and adenine). Transformed cells are first plated on media lacking L/T to select for cells successfully transformed with both plasmids. These co-transformed cells are then grown in the −L/T selective media and plated on the stringent −L/T/H/A media; growth on −L/T/H/A media indicates positive interaction of bait and prey proteins.

### Data and reagent availability

Strains and plasmids are available upon request.

## RESULTS

### Reducing Erk levels suppresses the abnormal locomotion and neuronal dysfunction exhibited by *sax-7* null animals

We previously determined that knocking out *ksr-1* suppresses the abnormal locomotion exhibited by *sax-7* null animals (Moseley-Alldredge *et al*. 2022). The KSR-1 protein acts as a scaffold for the core kinase components of the MAPK cascade to facilitate efficient activation of ERK (Nguyen *et al*. 2002; Roy *et al*. 2002; Brennan *et al*. 2011; Sundaram 2013; Frodyma *et al*. 2017). This result suggests that reduced levels of activated ERK underlie the rescue in *sax-7; ksr-1* locomtion abnormalities. We thus reasoned reduced levels of ERK, encoded by the *mpk-1* gene, would similarly suppress *sax-7* mutant locomotion phenotypes. To test this possibility, we crossed into *sax-7* animals the *mpk-1(ga117)* null allele (Lackner *et al*. 1994) and examined the impact of the loss of MPK-1/ERK on three locomotion phenotypes displayed by *sax-7* null animals.

The first locomotory characteristic we examined was the ability for animals to disperse when crawling on solid medium. Tracing the movement of individual animals, we can observe the overall radial displacement for each animal. Wild-type animals tend to make long and directed runs that radiate from the point of origin, which is set as the middle of the graph (0,0 coordinate) shown in figure 1A; as such, wild-type animals show robust radial displacement within a one minute interval. The locomotory behavior of *sax-7* mutant animals is described best as loopy with exaggerated body bends and a tendency to meander locally. Consistent with this behavior, *sax-7* mutant animals tend to remain within the vicinity of the point of origin after the one-minute interval (Fig 1A). As a result, *sax-7* null animals exhibit poor radial displacement (Moseley-Alldredge *et al*. 2022). This phenotype is suppressed when *mpk-1* function is knocked out in *sax-7* null animals; indeed, *mpk-1; sax-7* animals generally make longer and more directed runs that radiate from the point of origin. *mpk-1* null animals, which do not exhibit apparent locomotory abnormalities, present a similar radial displacement as wild-type animals (Fig 1A) (Moseley-Alldredge *et al*. 2022).

**Figure 1:**
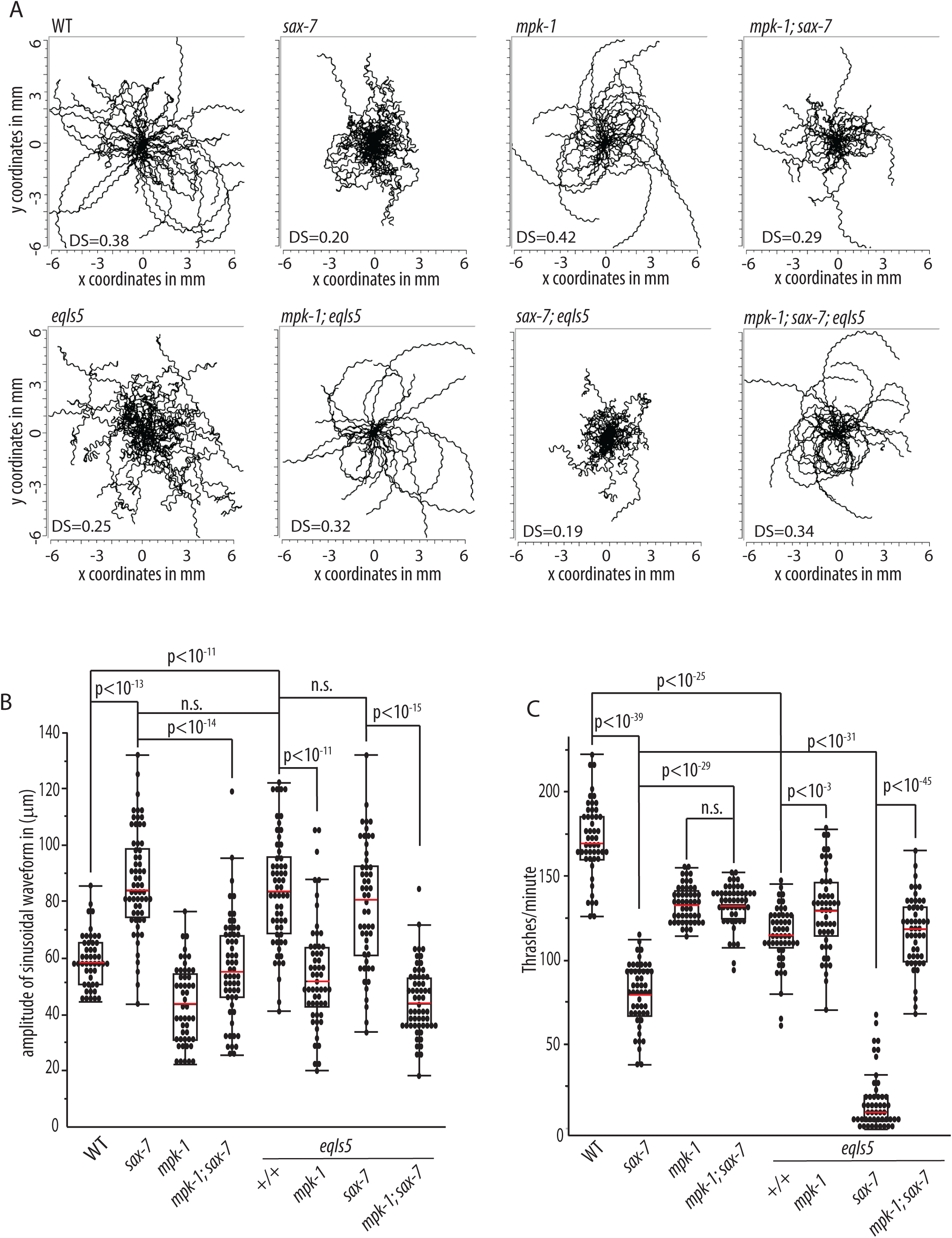
Loss of *mpk-1* function suppresses locomotory abnormalities exhibited by *sax-7* null animals as well as *eqIs5* animals overexpressing KSR-1 in cholinergic neurons. A. Graphs tracing the movements of 50 random animals crawling on agar media for 1 minute. Each line represents the track of an individual animal radiating from the point of origin in the center (0,0 coordinate). Directness Score (DS), a measurement of the average straightness of the worm trajectory, is provided for each strain. B. A box and whisker plot showing the average amplitude of the sinusoidal waveform of 50 random animals per strain. Loss of *mpk-1* corrects the exaggerated waveform of *sax-7* animals. C. A box and whisker plot showing the average swim rate (thrashes per minute) for 50 animals per strain. For each box and whisker plot, the middle red line in the box is the mean, the bottom and top of the box are the 25^th^ and 75^th^ quartiles, and the whiskers extend to the 1.5 interquartile range from the top and bottom of the box. n = 50 per strain, p-values are shown, n.s., not significant, two-way ANOVA with Bonferroni’s *post hoc* test.

Radial displacement can be affected by different behaviors, including crawling speed or the motivation to move. To uncouple local meandering movements from such behaviors, we measured the straightness of an animal’s trajectory by calculating a directional score (DS): the ratio of the average radial displacement to the average total distance travelled. Whereas the average DS for the wild-type strain is 0.38, the *sax-7* null strain has a DS of 0.2, consistent with the tendency for *sax-7* null animals to meander locally; *i.e.* reduced dispersal despite active crawling. On the other hand, the *mpk-1; sax-7* strain has an improved DS of 0.29 relative to the *sax-7* null strain, consistent with loss of *mpk-1* function rescuing the poor radial displacement exhibited by *sax-7* null animals.

The second characteristic we examined is the locomotory posture, which is reflected by the sinusoidal footprints left by animals crawling on solid agar medium. The tracks wild-type animals make appear as regular sinusoidal waveforms (Fig 1A) that have an average amplitude of 59μm (Fig 1B). On the other hand, the tracks *sax-7* mutant animals generate have a sinusoidal waveform that is irregular and an increased amplitude of 86μm, reflective of their loopy locomotory posture with exaggerated body bends. Loss of *mpk-1* rescues this phenotype so that the sinusoidal footprints *mpk-1; sax-7* animals make are more regular and have an average amplitude of 54μm, similar to wild-type tracks.

The third locomotory trait we examined is the ability for animals to swim (Fig 1C). *sax-7* mutant animals have reduced neuronal function that is readily apparent when placed in liquid (opperman *et al*. 2015; Moseley-Alldredge *et al*. 2022). *sax-7* null animals exhibit a decreased average swim rate of ~80 thrashes/minute, which is 47% the swim rate of wild-type animals at 152 thrashes/minute. Relative to the *sax-7* null strain, *mpk-1; sax-7* mutant animals display an improved swim rate of 131 thrashes/minute. When taken together, these results confirm our prediction that loss of *mpk-1* rescues the locomotory abnormalities exhibited by *sax-7* mutant animals.

### Elevated Erk signaling in cholinergic neurons induces similar locomotory behaviors observed in *sax-*7 null animals

How does loss of *mpk-1* function suppress *sax-7* locomotory abnormalities? One hypothesis is that *sax-7* and *mpk-1* function in parallel and opposing pathways to control coordinated locomotion. In the absence of *sax-7* function, MPK-1/ERK signaling is left unchallenged, causing uncoordinated locomotion. An alternative hypothesis is that *sax-7* acts in the same pathway as *mpk-1*, negatively regulating MPK-1/ERK activation or function. Thus loss of *sax-7* function would result in increased levels of activated MPK-1/ERK that lead to the observed locomotory abnormalities. Regardless of how *sax-7* opposes Erk signaling, both hypotheses predict that elevated MPK-1/ERK signaling in wild-type animals would phenocopy *sax-7* null animals. Because KSR-1 facilitates efficient activation of the MAPK module of kinases, we reasoned that KSR-1 overexpression would lead to increased levels of activated MPK-1 and thus elevated MPK-1/ERK signaling (Levchenko *et al*. 2000; Kortum and Lewis 2004). We had previously determined that KSR-1 acts in cholinergic neurons to influence *sax-7-*mediated locomotion (Moseley-Alldredge *et al*. 2022). To test our prediction, we generated *eqIs5*, an integrated multicopy transgene that drives KSR-1 overexpression in cholinergic neurons using the promoter for the *unc-17* acetylcholine transporter (Alfonso *et al*. 1994).

*eqIs5* animals exhibit a loopy locomotory behavior that is strikingly similar to that of *sax-7* mutant animals (Fig 1). Their reduced radial displacement is evident by their tendency to remain within the vicinity of the original placement site and a reduced DS of 0.25 (Fig 1A). Reflective of their loopy posture, their sinusoidal footprint is irregular and has an increased amplitude of 83μm (Fig 1B). *eqIs5* animals also show apparent neuronal deficits with a reduced swim rate of 113.7 thrashes/minute, which is 67% of the wild-type swim rate (Fig 1C). The fact that *sax-7* null animals have a lower swim rate than *eqIs5* animals is not surprising as *sax-7* function is required in additional neurons besides acetylcholine neurons for full neuronal function (Moseley-Alldredge *et al*. 2022). Importantly, all three *eqIs5-*induced locomotory abnormalities are suppressed by the loss of *mpk-1* function, consistent with our theories that 1) KSR-1 overexpression results in elevated MPK-1/ERK signaling and 2) elevated MPK-1/Erk signaling contributes to the locomotory phenotypes exhibited by *sax-7* null animals. Indeed, relative to *eqIs5* animals, *mpk-1; eqIs5* double mutant animals have significantly improved radial displacement with longer and more vectorial runs and a recovered DS value of 0.32. Their sinusoidal footprint has a 54μm amplitude, similar to wild-type tracks. Lastly, the *mpk-1; eqIs5* swim rate of ~130 thrashes/min is restored to the level seen in *mpk-1* null animals. While *mpk-1* null animals do not display obvious locomotory abnormalities when crawling on solid media, the modest reduction in their swim rate relative to wild-type animals indicates some neuronal deficits.

We next assessed how loss of *sax-7* might influence the effects of *eqIs5* (Fig 1). On a gross level, the locomotory behavior of *sax-7; eqIs5* double mutant animals appeared slightly more pronounced than *eqIs5* animals. However, the radial displacement exhibited by *sax-7; eqIs5* animals and the DS value of 0.19 is virtually identical to that of *sax-7* single mutant animals. Moreover, the 78.7μm amplitude of *sax-7; eqIs5* sinusoidal footprint is not significantly different from that of either *sax-7* or *eqIs5* single mutant animals. However, relative to *sax-7* and *eqIs5* single mutant animals, *sax-7; eqIs5* animals exhibit a synergistically reduced swim rate of 21 thrashes/minute, which may explain the seemingly more pronounced abnormal locomotion. Importantly, knocking out *mpk-1* function in *sax-7; eqIs5* animals dramatically suppresses all three traits. Taken together, these results strongly support the idea that elevated MPK-1/ERK signaling in cholinergic neurons underlies the locomotory posture and neuronal deficits exhibited by *sax-7* null animals.

### Elevated Erk signaling synergizes with loss of *sax-7* in non-neuronal tissues

A key upstream activator of the Erk signaling pathway is the Ras GTPase, encoded by *let-60* in *C. elegans*. The *let-60*(*n1046*) allele is a gain-of-function G13E missense allele (*let-60(gf)* hereafter) that results in elevated MPK-1/ERK signaling (Beitel *et al*. 1990). Thus, the *let-60(gf)* allele provides us an additional approach to assess whether elevated MPK-1/ERK activity can phenocopy *sax-7* null animals. A gross examination of *let-60(gf)* animals did not show obvious locomotory phenotypes. One possible reason is *let-60* does not play a role in regulating locomotion, as previously shown (Coleman *et al*. 2018). An alternative possibility is that the activated Erk levels in *let-60(gf)* animals may not be at a threshold required for a visible neuronal phenotype. Because *eqIs5* further decreases the already reduced swim rates exhibited by *sax-7* null animals, we reasoned we might similarly observe enhanced phenotypes when *let-60(gf)* is introduced into *sax-7* null animals.

In fact, we did observe synergy between *sax-7* and *let-*60 that was manifested in three distinct and unexpected non-neuronal phenotypes, revealing previously uncharacterized roles for *sax-7*. First, we detected in many *sax-7 let-60(gf)* animals a prominent protrusion at the excretory pore (Fig 2A). This protuberance is absent in *sax-7* null animals and modest in *let-60(gf)* animals. The presence and severity of this excretory pore abnormality in two different *sax-7 let-60(gf)* strains using the *sax-7* alleles, *eq1* and *eq22* (Fig 2Aiii), indicates that enhancement of this *let-60(gf)* phenotype is dependent on impaired *sax-7* function and is not allele specific.

**Figure 2:**
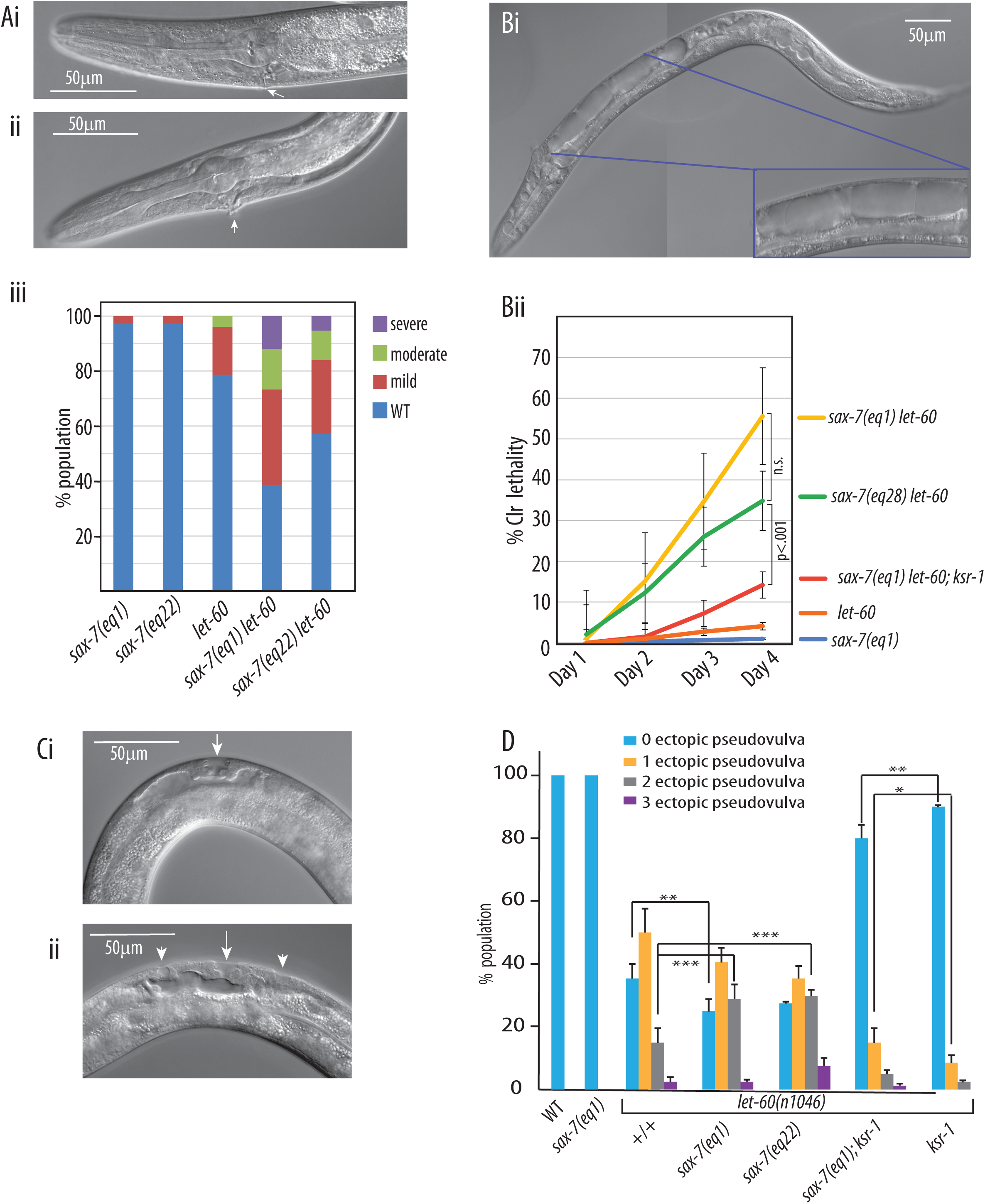
Loss of *sax-7* function synergizes with elevated Ras activity outside the nervous system, revealing non-neuronal roles for SAX-7. (A) Synergistic enhancement of excretory pore abnormalities are observed in *sax-7 let-60(gf)* animals. DIC microscopy images of a wild-type excretory pore in *sax-*7(*eq1*) animals (Ai, arrow) and an abnormal excretory pore with a protuberance in sax*-7; let-60(gf)* animals (Aii, arrow). Depending on how far the protrusion extends, we have categorized the protrusion as mild, moderate (Aii, arrow), and severe, which is notable for apparent cellular debris spilling out of the pore. (Aiii) A graph showing the distribution of mild, moderate, and severe excretory pore abnormality; n = 75 per strain. (B) Synergistic enhancement of Clr lethality is observed in *sax-7 let-60(gf)* animals, as shown in the DIC micrograph (Bi; see insert for a magnified image of the apparent fluid build-up underlying the Clr phenotype). This Clr phenotype is progressive with lethality typically ensuing shortly after the animals adopt the Clr appearance (Bii). Error bars show standard error of the mean (SEM) of 3 sample sets where n = 100 animals for each set. n.s. non significant, p values as shown with two-way ANOVA with Bonferroni’s *post hoc* test. (C) Synergistic vulval hyperinduction is observed in *sax-7 let-60(gf)*. DIC micrographs of the developing vulva in L4-staged larvae show a wild-type developing vulva in a *sax-7(eq1)* animal, as indicated by the characteristic single “Christmas tree” invagination (Ci, arrow). In *sax-7; let-60(gf)* mutant animals (Cii), adjacent to the developing vulva (arrow) are additional smaller invaginations (arrowheads) that are characteristic of ectopic vulval inductions. (D) Quantification of animals with a wild-type vulval development and those with additional pseudovulvae. Relative to *let-60(gf)* animals, a significantly higher percentage of *sax-7; let-60(gf)* animals exhibit pseudovulvae. Moreover, there are more animals with 2 ectopic invaginations, showing a more severe Muv phenotype in *sax-7; let-60(gf)* animals, relative to *let-60(gf)* animals. Error bars show SEM of 3 sample sets where n = 50 animals for each set; p-values: *p < 0.05, **p < 0.01, ***p < 0.005, n.s., not significant, two-way ANOVA with Bonferroni’s *post hoc* test.

Second, we discerned in many *sax-7 let-60(gf)* double mutant adult animals a **Cl**ea**r** (Clr), more transparent body that is not exhibited in *sax-7* null animals and minimally in the *let-60(gf)* strain (Fig 2B). Clr *sax-7 let-60(gf)* animals are first detected two days post-hatching, with numbers increasing on the third day when most animals are L4-staged larvae. Many of the Clr animals become immobile, dying a day or two after becoming Clr. This progressive Clr lethality is observed in two *sax-7 let-60(gf)* strains using the *sax-7* null alleles, *eq1* and *eq28,* indicating the genetic interaction with *let-60* is not allele specific but rather dependent on the loss of *sax-7*. Ras can activate other signaling pathways in addition to the MAPK-ERK pathway (Sundaram 2013; KIEL *et al*. 2021; Wu and Reiner 2022; Smith 2023). To determine whether this interaction between *sax-7* and *let-60(gf)* is dependent on MPK-1/ERK signaling, we tested whether loss of *ksr-1* function suppresses the *sax-7 let-60(gf)* Clr lethality. *sax-7 let-60; ksr-1* triple mutant animals exhibited significantly reduced Clr lethality, consistent with the notion that this phenotype is a consequence of elevated MPK-1/Erk signaling.

Third, we observed many *sax-7 let-60(gf)* adult animals displaying the **Mu**ltiple **v**ulva (Muv) phenotype. While the Muv phenotype is characteristic of *let-60(gf)* animals (Beitel *et al*. 1990), the *sax-7 let-60(gf)* strain appears to have a more severe Muv phenotype, based on the number of adult animals displaying an increased number of pseudovulvae protuberances flanking the vulva along the ventral midline. To confirm this perception, we examined the developing vulva in *sax-7 let-60(gf)* animals in more detail using Differential Interference Contrast (DIC) microscopy.

The developing vulva is best examined in the L4-staged larva when it appears as a single invagination with a characteristic “Christmas tree” morphology, mid-point along the anterior-posterior axis on the ventral midline of the animal (Fig 2Ci, arrow). Vulval development starts in the L3 larval stage with the anchor cell (AC), located in the gonad, secreting LIN-3/EGF (epidermal growth factor) that diffuses as a gradient to the six vulval precursor cells (VPC) located along the ventral midline. Activation of the LET-23 EGF receptors on the VPCs triggers the Ras-MAPK signaling cascade to induce the 1° vulval fate in the VPC closest to the AC (Fig 3A). The induced VPC subsequently induces the flanking VPCs to assume the 2° vulval fate via the LIN-12/Notch signaling pathway; together, the three VPCs and their descendants contribute the 22 cells that make up the vulva (Sternberg 2005; Sundaram 2013; Gauthier and Rocheleau 2017; Shin and Reiner 2018). With excessive Ras-MAPK signaling, as in *let-60(gf)* animals, 1° vulval inductions can also occur in additional VPCs independent of EGF receptor activation, leading to ectopic smaller invaginations (Fig 2Cii, arrowheads) that flank the developing vulva (Fig 2Cii, arrow); by adulthood, these small invaginations often develop into protruding pseudovulvae that are visible under the dissecting microscope.

**Fig 3:**
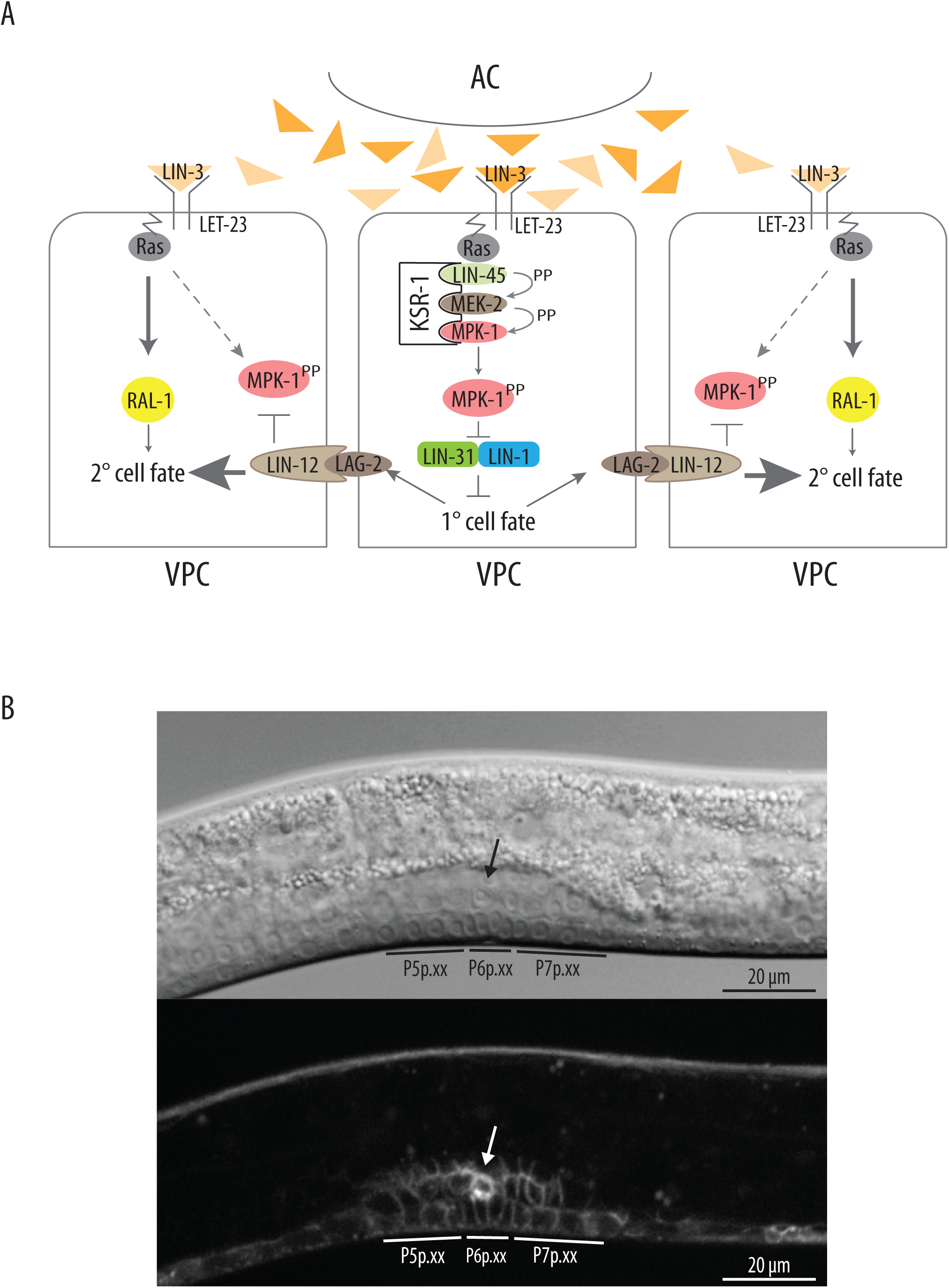
SAX-7 is localized to the plasma membrane of the Anchor Cell and Vulval Precursor Cells. (A) A schematic of the EGF-Ras-Erk signaling pathway necessary for vulval induction. The LET-23/EGF receptors, when activated by the LIN-3/EGF ligand secreted from the AC, trigger the RAS-ERK signaling pathway that promote 1° vulval induction in the VPC closest to the AC. Activated MPK-1/ERK phosphorylates the complexed transcription factors, LIN-31 and LIN-1, resulting in their dissociation and thus permitting each transcription factor to regulate gene expression, thereby promoting 1° vulval cell fate. Flanking VPCs subsequently undergo 2° vulval induction, triggered primarily by LIN-12/Notch signaling and to a lesser degree, RAL signaling. (B) DIC (upper panel) and fluorescence micrographs (lower panel) show that endogenous SAX-7::mCherry fusion protein is present on the plasma membrane of the AC (arrow), along with other cells in the gonad, as well as the granddaughters of the induced VPCs (P5p, P6p, P7p).

To determine whether loss of *sax-7* function enhances the Muv phenotype in the *let-60(gf)* strain, we examined two *sax-7 let-60(gf)* strains using the *sax-7* alleles, *eq1* and *eq22;* control strains included in this analysis are wild-type, *let-60(gf),* and *sax-7(eq1)*. We did not observe any vulval abnormalities in wild-type and *sax-7* null L4-staged larvae. In contrast and as expected, we observed in the *let-60(gf)* control strain ectopic invaginations that are characteristic of excessive 1° vulval induction (Fig 2D). Specifically, 35% of *let-60(gf)* displayed a single developing vulva with no ectopic invagination, typical of wild type vulval development, while ~ 50% showed one additional ectopic invagination and ~15%, two ectopic invaginations (Fig 2D). On the other hand, only 25% of *sax-7*(*eq1*) *let-60(gf)* and 27% of *sax-7(eq22) let-60(gf)* animals displayed normal vulval development, with the remaining percentage displaying a Muv phenotype. Notably, 28% of *sax-7*(*eq1*) *let-60(gf)* and 30% of *sax-7(eq22) let-60(gf)* animals exhibited two ectopic invaginations (Fig 2D), a significant increase relative to *let-60(gf)* single mutant animals with two ectopic invaginations. Taken together, these findings support our first impressions that the Muv phenotype is stronger in the *sax-7 let-60(gf)* strain, as compared to the *let-60(gf)* strain. Importantly, knocking out *ksr-1* function dramatically suppresses the Muv phenotype in both *sax-7 let-60(gf)* and *let-60(gf)* animals (Fig 2D), consistent with elevated MPK-1/Erk signaling as promoting the Muv phenotype. The modest but significant increase in the percentage of Muv animals in the *sax-7 let-60(gf); ksr-1* strain relative to the *let-60(gf); ksr-1* strain further highlights a role for *sax-7* in vulval development.

### SAX-7 is expressed in both the vulval precursor cells and the anchor cell

The findings presented thus far are consistent with SAX-7 acting as an antagonistic modulator of MPK-1/ERK signaling. With the genetic interaction between *sax-7* and *mpk-1* impacting such diverse tissues and taking parsimony into consideration, we hypothesize that *sax-7* likely functions in same pathway as *mpk-1,* rather than acting in parallel pathways. Because vulval development is such a well-established paradigm for MPK-1/ERK signaling, we reasoned it would be advantageous to use the developing vulva to test our hypothesis and dissect how SAX-7 modulates MPK-1 function. To start, we assessed whether *sax-7* is expressed in cells that are essential for vulval development, such as the VPCs and/or the AC. We had previously reported widespread *sax-7* expression, based on immunofluorescence studies with an anti-SAX-7 polyclonal antibody; SAX-7 was detected in virtually all cells as early as the two-cell stage embryo with high expression levels in the nervous system, body-wall muscles, the hypodermis, seam cells, and the intestine (Chen *et al*. 2001; Zhou *et al*. 2008). This expression was confirmed using the *sax-7(eq23)* allele, which allows detection of endogenous SAX-7::mCherry expression (Moseley-Alldredge *et al*. 2022). Using *sax-7(eq23)*, we detected SAX-7::mCherry on the plasma membrane of the VPCs and their descendents as well as the AC, along with other cells in the gonad (Fig 3B).

### SAX-7 acts in the Vulval Precursor Cells to antagonize LET-23-mediated vulval induction

The enhanced Muv phenotype in *sax-7 let-60(gf)* animals suggests that loss of *sax-7* leads to increased vulval induction. If this notion is correct, we predict that loss of *sax-7* function would suppress the **Vul**valess (Vul) phenotype that is characteristic of animals with reduced levels of MPK-1/ERK signaling, such as animals with a defective EGF receptor, encoded by the *let-23* gene. Because *let-23* null animals die prior to vulval induction, we used the *let-23(sy1)* hypomorphic allele in our analysis. Consistent with previous analysis (Aroian and Sternberg 1991), *let-23* hermaphrodites are viable and Vul. Specifically, 85% of L4-staged animals lacked any invagination, which is typical of deficient vulval induction, harboring visible uninduced daughters of the six VPCs (arrowheads in Fig 4Ai, B). The remaining 15% of *let-23* animals exhibited invaginations indicative of some vulval induction. Most of these invaginations are small, typical of a partial vulval induction (black arrow in Fig 4Aii), while a handful of animals exhibit the characteristic “Christmas tree” morphology that is representative of a wild-type developing vulva (white arrow in Fig 4Aiii (Sternberg 2005; Gauthier and Rocheleau 2017)). In contrast, 65% of *let-23; sax-7(eq1)* animals are Vul with the remaining 35% showing a combination of incomplete and complete vulval induction (Fig 4B). This suppression of the *let-23* Vul phenotype by loss of *sax-7* function is in agreement with our conjecture of SAX-7 opposing signals that promote vulval induction.

**Fig 4:**
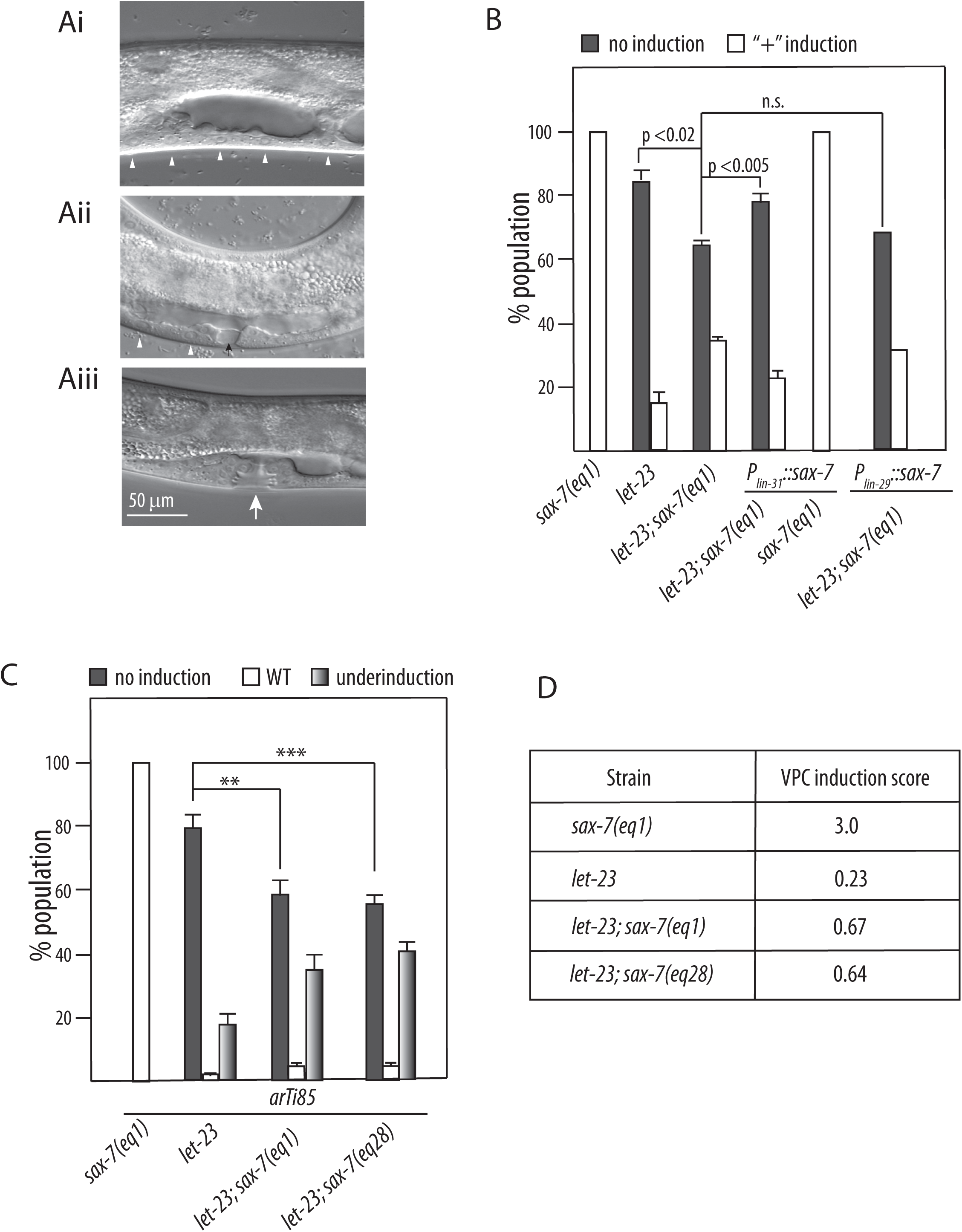
Loss of SAX-7 rescues vulval induction in *let-23* animals. (A) DIC micrographs of hypomorphic *let-23* L4-staged larvae illustrates the spectrum of vulval induction in this strain, ranging from absent (Ai, arrowheads point to uninduced VPC daughter cells) to partial (Aii, black arrow) and complete (Aiii, white arrow) with a wild-type “Christmas tree” morphology. (B) Examination of the developing vulva via DIC microscopy revealed that loss of *sax-7* function suppresses the absence of vulval induction exhibited in *let-23* L4-staged larvae. This suppression is reversed in *let-23; sax-7* animals with targeted *sax-7* expression in the VPCs using P*_lin-31_*::*sax-7* transgenes but not in the AC using P*_lin-29_*::sax-7 transgenes. (C) Vulval induction examined with the aid of a VPC-targeted fluorescent reporter expressed from the *arTi85* transgene. This analysis confirmed the finding that loss of *sax-7* function (with null alleles, *eq1* and *eq28*) suppresses the absence of vulval induction in *let-23* animals so that there is a significant increase in the number of animals with partial vulval induction. (D) The fluorescent reporter allowed us to calculate the VPC induction score. As described in the body of the paper, wild-type animals have a vulval induction score of “3” because induction of three VPCs is necessary for a wild-type vulva to form. Here, the vulval induction score is significantly higher in *let-23; sax-7* animals than in *let-23* animals. Error bars show SEM of 3 sample sets where n = 50 animals for each set; p-values as shown with one-way ANOVA with Bonferroni’s *post hoc* test.

To better discern the extent of vulval induction, we examined the VPCs in both *let-23* and *let-23; sax-7* strains with the aid of a VPC-specific mCherry reporter expressed by the *arTi85* transgene (De La Cova *et al*. 2017). Consistent with our assessment based on the morphology of the developing vulva, the *let-23; sax-7* strain showed a higher level of vulval induction as compared to the *let-23* single mutant strain (Fig 4C). We observed an absence of vulval induction in ~60% of *let-23; sax-7(eq1)* animals. Moreover, ~35% of *let-23; sax-7(eq1)* animals exhibited partial vulval induction as opposed to 18% of *let-23* mutant animals while 5% of *let-23; sax-7(eq1)* animals displayed complete vulval induction with only 1% *let-23* animals doing so. This distribution of vulval induction was also observed in another *let-23; sax-7* strain using the *sax-7(eq28)* null allele. Here, 55% of *let-23(sy1); sax-7(eq28)* animals lacked vulval induction while 41% showed incomplete vulval induction and 4%, complete vulval induction. The VPC-specific mCherry reporter also allowed us to calculate the VPC induction score, a quantification of vulval induction. In wild-type animals, the VPC induction score is three because three of the six VPCs and their daughters are induced to contribute the 22 cells that form the vulva (Sternberg 2005; Gauthier and Rocheleau 2017). The *sax-7(eq1)* strain has a VPC induction score of three, consistent with our data based on vulva morphology that vulval development is wild-type in *sax-7* null animals (Fig 4C, D). On the other hand, the VPC induction score for the *let-23(sy1)* strain is 0.23, indicative of reduced vulval induction. Both the *let-23(sy1); sax-7(eq1)* and *let-23(sy1); sax-7(eq28)* strains showed an higher VPC induction score of 0.67 and 0.64, respectively, in agreement with the notion that loss of *sax-7* results in increased vulval induction. The results presented here are remarkably consistent, regardless of whether we used vulval morphology or the VPC-specific fluorescent reporter to assess vulval development. For convenience, we thus relied on vulval morphology as the basis of our subsequence analyses.

We next sought to identify the cells in which *sax-7* functions in vulval development. Based on our hypothesis that SAX-7 and MPK-1 functions in the same pathway, we reasoned that *sax-7* is likely to function in the VPCs where MPK-1 functions to promote 1° vulval cell fate. We employed the *lin-31* promoter to drive *sax-7* expression in the VPCs in *let-23; sax-7* mutant animals (Miller *et al*. 1996b). If *sax-7* functions in the VPCs, we expect VPC-directed *sax-7* expression to reverse the suppression of the Vul phenotype in *let-23; sax-7* animals. Consistent with this prediction, we observed a significant increase in the percentage of Vul hermaphrodites in *let-23; sax-7* animals carrying *P_lin-31_*::*sax-7* extrachromosomal array as compared to *let-23; sax-7* animals (Fig 4B). While 65% *let-23; sax-7* mutant animals are Vul, 77% of *let-23; sax-7* mutant animals with VPC-directed *sax-7* expression are Vul. As control, we show that the *P_lin-31_*::*sax-7* extrachromosomal array does not impact vulval development in *sax-7* null animals. Thus, this result indicates that *sax-7* expression in the VPCs is sufficient for *sax-7* to modulate vulval development. In contrast, targeted *sax-7* expression in the anchor cell using the *P_lin-29_*::*sax-7* extrachromosomal array did not reverse the suppression of the vulvaless phenotype in the *let-23; sax-7* strain.

### The SAX-7 role in vulval development is dependent on the pro 1° vulval fate MPK-1/Erk pathway

We have thus far examined how *sax-7* influences the signaling components upstream of the Erk signaling pathway. Although *let-23* and *let-60* are necessary in Erk-mediated pro-1° vulval fate signaling pathway, they also function in a secondary pro-2° vulval induction pathway mediated by the Ral GTPase (Zand *et al*. 2011; Shin and Reiner 2018) (see Fig 3A). To assess the interplay between *sax-7* and specifically the Erk signaling pathway, we examined how loss of *sax-7* influences vulval development in animals with defective RAF, encoded by *lin-45*. Because loss of *lin-45* function results in early larval lethality prior to vulval induction, we used the *lin-45(sy96)* hypormophic allele in our analysis. We observed an absence of vulval induction in ~30% of *lin-45* animals with the remaining 70% of animals showing positive vulval induction that range from partial to complete (Fig 5A). On the other hand, only 18% of *sax-7(eq22) lin-45* animals lacked signs of vulval induction; the remaining 82% of *sax-7(eq22) lin-45* animals showed positive vulval induction, with most animals exhibiting partial induction and the remaining, complete induction.

**Fig 5:**
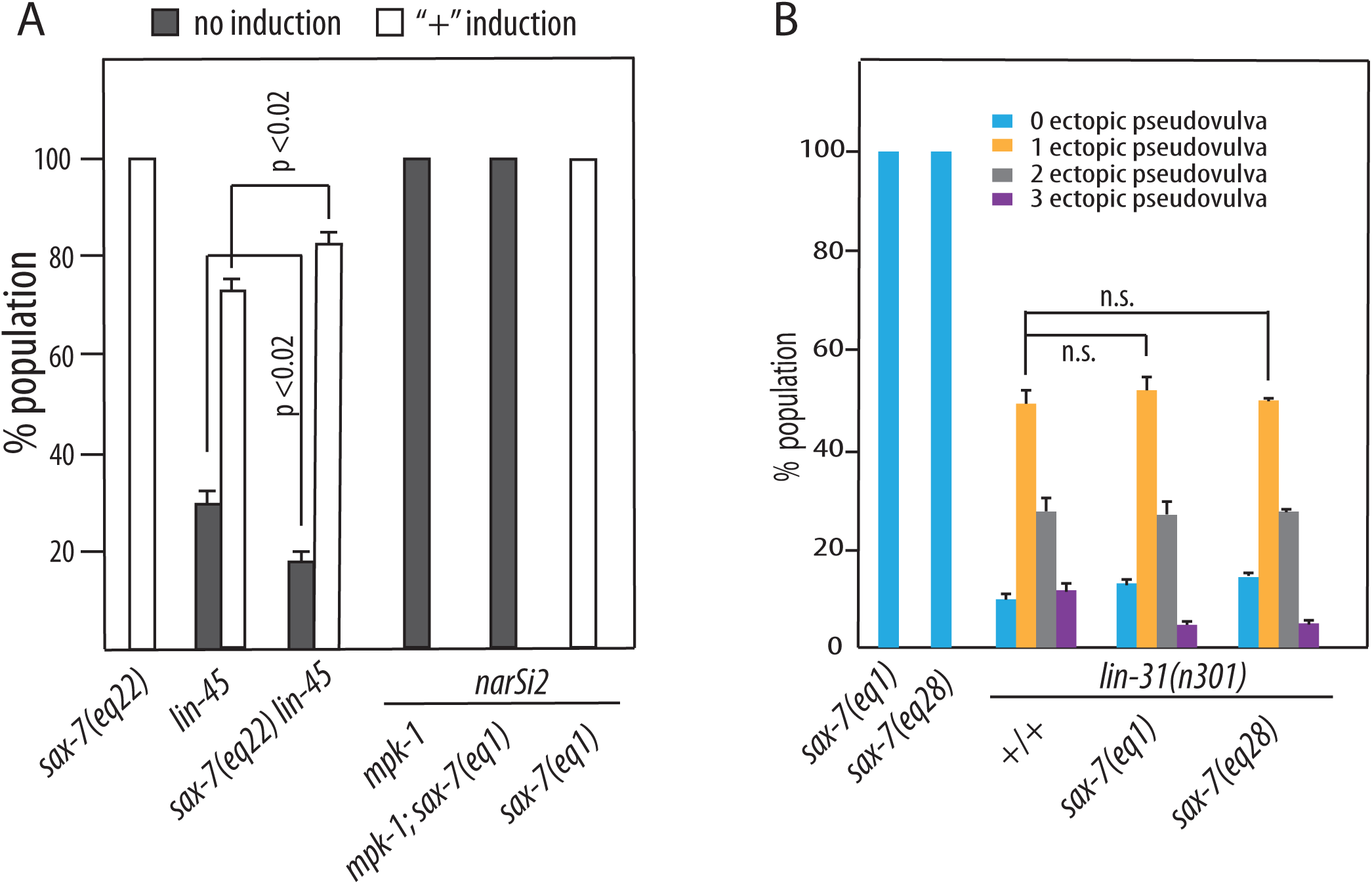
SAX-7 acts in the MPK-1/ERK signaling pathway to regulate vulval development. (A) Loss of *sax-7* can suppress the lack of vulval induction in *lin-45* hypomorphic animals with reduced RAF function, but not in *mpk-1* animals that lack somatic ERK function; here, the *narIs2* integrated transgene provides the germline-specific *mpk-1* isoform to rescue the sterility exhibited by *mpk-1* null animals. This result indicates the role *sax-7* plays in vulval development is dependent on the MPK-1/ERK pathway. (B) Consistent with *sax-7* acting in the MPK-1/ERK pathway, loss of *sax-7* function does not enhance *lin-31* null Muv phenotype. Error bars show SEM of 3 sample sets where n = 50 animals for each set; p-values as shown with two-way ANOVA with Bonferroni’s *post hoc* test.

MPK-1/ERK is necessary for vulval formation. As previously demonstrated (Lackner *et al*. 1994), we observed the Vul phenotype in 100% of *mpk-1* animals (Fig 5A). We also observed a lack of vulval induction in 100% of *mpk-1; sax-7(eq1)* animals, indicating that the role *sax-7* plays in vulval development is dependent on MPK-1. To validate this notion further, we next examined how *sax-7* interacts with genes encoding transcription factors controlled by MPK-1/Erk in vulval development.

The LIN-31 and LIN-1 transcription factors are MPK-1 substrates in the pro 1° vulval pathway (Jacobs *et al*. 1998; Tan *et al*. 1998; Sternberg 2005; Sundaram 2013). Both transcription factors form a complex, disassociating when phosphorylated by MPK-1 to trigger 1° vulval induction (Fig 3A). Thus loss of either transcription factor results in vulval hyperinduction, even in the absence of MPK-1 signaling, thus leading to a Muv phenotype. Indeed, only 10% of *lin-31* null L4-staged larvae exhibited a single invagination typical of a wild-type developing vulva while 90% exhibited ectopic invaginations in addition to the developing vulva. Specifically ~50% of *lin-31* animals displayed one ectopic invagination, ~28% showed two ectopic invaginations, and ~12%, three ectopic invaginations (Fig 5B). If *sax-7* acts in the same pathway as MPK-1/ERK signaling, then loss of *sax-7* function should not affect the Muv phenotype in *lin-31* null animals. In contrast, if *sax-7* acts in a parallel pathway as *mpk-1,* then loss of *sax-7* function in *lin-31* null animals would increase the severity of the Muv phenotype. Relative to the *lin-31* single mutant strain, we did not observe a significant difference in either of the *lin-31; sax-7* strains (Fig 5B). These results are consistent with *sax-7* functioning in the *mpk-1* pathway.

### *gid-1* functions in the same pathway as *sax-7* to negatively modulate MPK-1/Erk signaling

How does SAX-7 modulate MPK-1/Erk signaling? One possibility is SAX-7 impedes activated Erk, perhaps limiting its availability through sequestration or promoting its downregulation via a MAPK phosphatase (Roskoski 2012; Sundaram 2013; Kidger and Keyse 2016). An alternative possibility is that SAX-7 hinders the activation of the Erk cascade. A previous study identified Ran-binding protein in the microtubule-organizing center (RanBPM, aka RanBP9 in mammals) as molecularly interacting with the mammalian L1CAM protein, linking L1CAM to the Erk signaling pathway to control neurite outgrowth (Cheng *et al*. 2005). RanBp9 is a ubiquitous scaffold protein that is highly conserved in metazoans. Despite its name, RanBP9 has weak, if any, ability to bind Ran. However, with its multiple protein-protein interacting domains, including SPRY (spore lysis A and ryanodine receptor), LisH (lissencephaly type-1-like homology), CTLH (carboxy terminal to LisH), and CRA (CT11-RanBPM), interactions with diverse proteins link RanBP9 to multiple intracellular signaling pathways (Salemi *et al*. 2017; Das *et al*. 2018). In fact, independent studies revealed RanBP9 as an integral component of the multi-subunit CTLH (C-terminal to LisH) E3 ubiquitin ligase complex, which targets substrate proteins for degradation. Interestingly, one such target substrate is c-RAF (Atabakhsh and Schild-Poulter 2012; Mctavish *et al*. 2019). Based on these studies, we hypothesized that the *C. elegans* RanBP9, encoded by the *gid-1* gene (Shaye and Greenwald 2011; Kim *et al*. 2018), might similarly link SAX-7 to the MPK-1/ERK signaling pathway to control LIN-45/Raf levels as part of the CTHL E3 ubiquitin ligase complex; in this way, GID-1 would mediate the antagonistic action of SAX-7 on ERK activation (Fig 6A).

**Fig 6:**
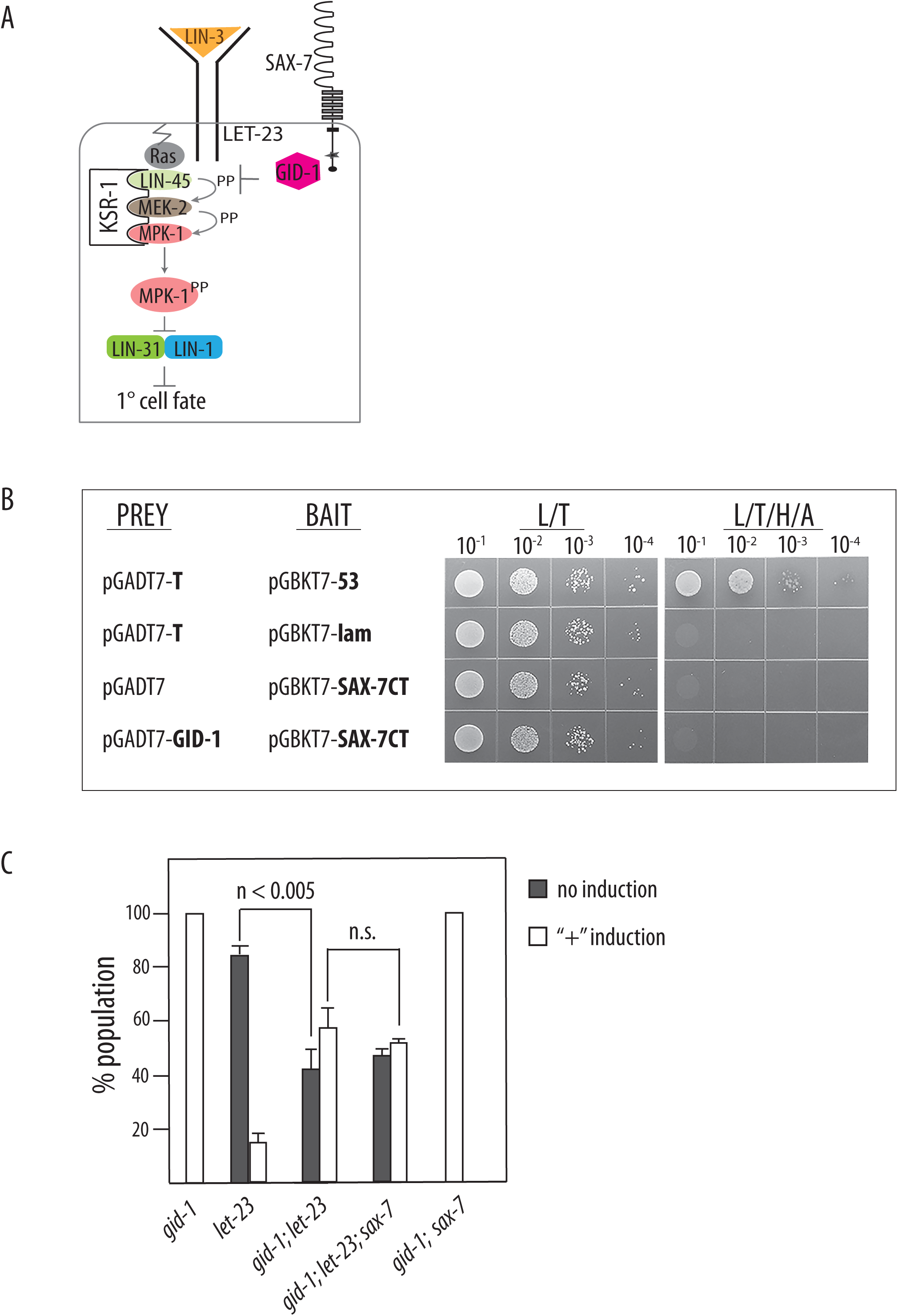
The RanBPM orthologue, GID-1, functions in the same pathway as SAX-7 to modulate vulval development. (A) A model of SAX-7 modulating MPK-1/ERK signaling via GID-1, a RanBP9 orthologue that molecularly interacts with mammalian L1CAM. RanBP9 acts in the CHTL E3 ubiquitin ligase complex demonstrated to target cRAF for protein degradation. (B) SAX-7 and GID-1 do not molecularly interact in a yeast-two-hybrid assay. (C) Similar to *sax-7*, loss of *gid-1* function also suppresses the defective vulval induction observed in *let-23* animals. Moreover, there is no significant difference in vulval induction levels between *gid-1; let-23* double and *gid-1; let-23; sax-7* triple mutant animals, consistent with *gid-1* and *sax-7* functioning in the same genetic pathway.

To test our hypothesis, we first determined whether SAX-7 and GID-1 molecularly interact with each other. RanBPM was previously shown in a yeast two hybrid assay to molecularly interact with the mammalian L1CAM cytoplasmic tail, which has well-established protein-protein interacting domains, including the FERM-binding motif and ankyrin-binding motif. The interacting sites were narrowed down to the last 28 amino acids of L1CAM immediately following the ankyrin-binding motif; these 28 amino acids do not comprise any recognizable protein-interaction motif and the SPRY domain in RanBPM (Cheng *et al*. 2005). We had previously used the yeast-two-hybrid assay to identify a molecular interaction between the SAX-7 cytoplasmic tail (SAX-7CT) and UNC-44/ankyrin (Zhou *et al*. 2008). Here, we used the same SAX-7CT bait in a yeast two-hybrid assay to test for an interaction with full-length GID-1 as prey. Briefly, yeast transformed with bait and prey were cultured on selective media lacking leucine and tryptophan (L/T); positive growth indicates successful co-transformation (Fig 6B). These cells were then cultured and plated onto selective media lacking 4 amino acids: leucine and tryptophan as well as histidine and adenine (L/T/H/A). If the bait and prey proteins interact, histidine and adenine expression would be induced, thus permitting growth of the yeast on this more stringent medium. As expected, yeast transformed with the positive control bait and prey proteins, T antigen and P53, showed positive growth on L/T/H/A media lacking the four amino acids while yeast transformed with the negative control bait and prey proteins, T antigen and Lam, did not. Yeast transformed with the SAX-7CT and GID-1 grew on L/T media but not on the more stringent L/T/H/A growth medium. This result indicates SAX-7CT and GID-1 do not interact via the yeast-two-hybrid assay.

Since the yeast-two-hybrid assay can produce false negative results, we further investigated the role of *gid-1* in vulval development genetically, using the *tm3703* allele in *gid-1* that was isolated from the Japanese National Bioresource Project. We confirmed that *tm3703* is a 705 bp out-of-frame deletion that is predicted to result in a premature stop and thus, a putative *gid-1* null allele (Sternberg *et al*. 2024). Similar to the *sax-7* null strain, animals homozygous for *gid-1(tm3703)* do not exhibit vulval abnormalities the allele suppresses the Vul phenotype exhibited by *let-23* mutant animals (Fig 6C). Indeed, vulval induction was absent in only 40% of *gid-1(tm3703); let-23* double mutant animals, as compared to over 80% of *let-23* animals; the remaining 60% of *gid-1(tm3703); let-23* animals exhibited positive vulval induction ranging from partial to complete induction. These results suggest that like *sax-7, gid-1* also functions antagonistically to the LET-23/RAS/ERK signaling in vulval development. This finding is in agreement with the RanBPM-containing CTLH E3 ligase complex in targeting RAF for degradation. We next assessed whether *gid-1* functions in the same pathway as *sax-7*. To do so, we compared vulval development in *gid-1; let-23; sax-7* triple and *gid-1; let-23* double mutant animals. If *sax-7* and *gid-1* function in the same pathway, we do not expect to see significant differences in vulval induction in both strains. On the other hand, if both genes function in parallel pathways, then the triple mutant strain is predicted to show a reduced percentage of Vul animals relative to each double mutant strain. Consistent with *gid-1* and *sax-7* functioning in the same pathway, we did not observe significant differences between both strains. In further support of this notion, *gid-1; sax-7* double mutant animals do not exhibit any vulval defects. These results identify GID-1 as a novel player in vulval development, acting together with SAX-7 to modulate MPK-1/ERK signaling.

## DISCUSSION

In this study, we examined the genetic interaction between *sax-7* and *mpk-1* to determine how both genes function in the nervous system to promote coordinated locomotion. In the process, we discovered this genetic interaction extends beyond the nervous system, revealing a previously-uncharacterized role for *sax-7* in vulval development. Our findings in both the nervous system and the developing vulva are consistent with SAX-7 as an antagonistic modulator of MPK-1/ERK signaling with *sax-7* acting in the same pathway as *mpk-1*. In determining how *sax-7* ensures proper vulval development, we uncovered *gid-1* as acting in the same genetic pathway as *sax-7* to oppose MPK-1/ERK function in vulval induction.

### SAX-7 in controlling locomotory posture and coordination

Our study starts with showing that loss of *mpk-1* function suppresses the locomotory abnormalities exhibited by *sax-7* null animals (Fig 1). This result suggests elevated and/or unopposed MPK-1/ERK signaling may contribute to the abnormalities. In support of this notion, we show that KSR-1 overexpression in cholinergic neurons with the *eqIs5* transgene phenocopies *sax-7* null animals. Not only is the DS and amplitude of the sinusoidal footprints of both *sax-7* null and *eqIs5* animals remarkably similar, but these phenotypes are not enhanced in *sax-7; eqIs5* double mutant animals. Furthermore, knocking out *mpk-1* function suppresses the locomotory abnormalities induced by *eqIs5*, consistent with KSR-1 overexpression resulting in elevated MPK-1/ERK signaling. In agreement with our findings, expression of an activated form of LIN-45/RAF in cholinergic neurons similarly results in loopy and uncoordinated locomotion (Coleman *et al*. 2018). Taken together, these findings support the idea that elevated and/or unopposed MPK-1/Erk signaling underlie the abnormal locomotion exhibited by *sax-7* null animals. The lack of enhancement in the loopy posture in *sax-7; eqIs5* animals is consistent with both *sax-7* and *mpk-1* function acting in the same pathway in controlling that locomotory process.

*sax-7* null animals also exhibit neuronal deficits as reflected in their decreased swim rate (Fig 1C). Intriguingly, *eqIs5* further exacerbates the *sax-7* null swim rate so that *sax-7; eqIs5* animals have synergistically lower swim rates. However, this reduced swim rate is suppressed by knocking out *mpk-1* function, suggesting two possibilities. First, MPK-1/ERK signaling in cholinergic neurons has an additional role that is non-overlapping from that of SAX-7 in controlling swimming capabilities. Indeed, *mpk-1* null animals exhibit reduced swim rates, indicating neuronal deficits. The second possibility is the combined level of MPK-1/ERK signaling in *sax-7; eqIs5* cholinergic neurons is at a collectively higher threshold that synergistically exacerbates neuronal function.

The loopy and exaggerated body bends exhibited by *sax-7* null animals are reminiscent of the phenotypes caused by the gain-of-function *egl-30* allele, which results in elevated heterotrimeric G protein Gq activity (Bastiani *et al*. 2003). Gq signaling in cholinergic neurons controls neurotransmission and locomotory posture via the the EGL-8 phospholipase C and RHO-1/RHOA small monomeric GTPase (Lackner *et al*. 1999; Miller *et al*. 1999; Mcmullan *et al*. 2006). Loss of *mpk-1* function suppresses the loopy posture caused by *egl-30(gf)* and *rho-1(gf)* gain-of-function alleles (Coleman *et al*. 2018). Additional studies will need to be conducted to determine how SAX-7 functions relative to the Gq, RHO-1, and EGL-8 signaling pathways. Further studies are also required to identify MPK-1 effectors in the nervous system that control locomotory posture and coordination. Candidate effectors should be similar to the identified mammalian neuronal ERK effectors, such as voltage-gated sodium, calcium, and potassium channels as well as postsynaptic scaffolding proteins, PSD-93 and PSD-95; these effectors regulate neuronal excitability, synaptic activity and plasticity (Adams *et al*. 2000; Martin *et al*. 2006; Schrader *et al*. 2006; Guo *et al*. 2012; Mao and Wang 2016).

### SAX-7 function in vulval development

A similar genetic interaction between *sax-7* and elevated Erk signaling was also observed outside the nervous system, manifesting as synthetic or enhanced phenotypes that include excretory pore structural abnormalities, progressive Clr lethality, and increased vulval induction. All three phenotypes are suggestive of defects in tissues with established *mpk-1* function. In fact, vulva development is a well-established paradigm for MPK-1/ERK signaling. Vulval induction is dependent on MPK-1, the absence of which results in vulvaless animals. Findings from our epistasis analyses are consistent with SAX-7 acting in opposition to MPK-1/ERK signaling. Indeed, *sax-7* null alleles can suppress the vulvaless phenotype caused by mutations in genes acting upstream of *mpk-1*. But, they do not increase vulval induction in animals with mutations in *mpk-1* or genes that function downstream, suggesting that *sax-7* and *mpk-1* function in the same pathway in vulval development, as in the nervous system.

How does *sax-7* function oppose MPK-1 function? One possible model is SAX-7 sequestering activated MPK-1, either directly or indirectly. Another model is SAX-7 targeting activated MPK-1 for downregulation or degradation. As SAX-7 does not harbor enzymatic domains, this model would require an intermediate protein(s); for example, MAPK phosphatases such as LIP-1, which dephosphorylates activated MPK-1 for downregulation during vulval development (Berset *et al*. 2001). Yet another model is SAX-7 negatively regulating the activation of MPK-1. Supporting this last model is our finding of GID-1/RanBP9 acting in the same genetic pathway as *sax-7* to oppose MPK-1-dependent vulval induction. In mammals, RanBP9 is a component of the multi-subunit CTLH E3 ligase complex, which targets proteins, including cRAF, for ubiquitin-mediated protein degradation (Mctavish *et al*. 2019). While RanBP9 molecularly interacts with mammalian L1CAM (Cheng *et al*. 2005), our yeast-two-hybrid assay did not reveal a physical interaction between SAX-7 and GID-1. In retrospect, this negative interaction is not surprising considering the last 28 amino acids in the L1CAM cytoplasmic tail that interacts with RanBP9 shares a 19% sequence identity with the same region of SAX-7; the L1CAM-interacting domain in RanBP9, also known as the SPRY domain, shares a 50% sequence identity (Madeira *et al*. 2024). Taken together, these results suggest that SAX-7 and GID-1 interact indirectly to negatively modulate MPK-1/ERK signaling. While additional studies will need to be conducted to determine whether *gid-1* similarly functions with *sax-7* in the nervous system, it is notable that RanBP9 binds multiple neuronal proteins, functions in brain development, and participates in the processing of β-amyloid that is strongly tied to Alzheimer’s Disease (Lakshmana *et al*. 2012; Palavicini *et al*. 2013; Palavicini *et al*. 2014; Salemi *et al*. 2017; Das *et al*. 2018).

### Other SAX-7 non-neuronal functions

The structural abnormalities at the excretory pore and progressive Clr lethality in *sax-7 let-60(gf)* animals suggest defects in the excretory system, the development of which relies on MPK-1 signaling. The excretory pore, along with excretory duct, canal cell, and the CAN neurons, comprise the excretory system, which is essential for fluid homeostasis. Laser ablation or genetic mutations impairing development of the excretory system results in fluid build-up in the pseudocoelomic cavity that causes a characteristic Clr appearance and death. Importantly, the decision to become the excretory duct versus pore cell hinges on MPK-1/ERK signaling with elevated signaling resulting in two duct cells and no pore cell and an absence of signaling resulting in two pore cells and no duct cell (Yochem *et al*. 1997; Abdus-Saboor *et al*. 2011; Sundaram 2013; Sundaram and Buechner 2016). In addition to the excretory system, elevated MPK-1/ERK signaling in the hypodermis due to dysregulated FGF (fibroblast growth factor) receptor activity also leads to fluid build-up in the pseudocoelomic cavity and lethality (Huang and Stern 2004; Rodriguez Torres *et al*. 2024). Previous studies established SAX-7 expression and function in the hypodermis (Wang *et al*. 2005; Zhou *et al*. 2008; Diaz-Balzac *et al*. 2016; Zou *et al*. 2016). Based on findings that SAX-7 negatively modulates MPK-1/ERK signaling, it is conceivable that *sax-7* null alleles synergize with *let-60(gf)* to produce a sufficiently elevated level of MPK-1/ERK signaling to cause fluid dysregulation and developmental deficits in the excretory system, thus resulting in the synthetic phenotypes in *sax-7 let-60(gf)* animals.

The Clr lethal phenotype displayed by *sax-7 let-60(gf)* animals occurs in a progressive fashion, starting first with a transparent body that becomes increasingly clearer with ensuing immobility and finally, death. This progressive characteristic is consistent with fluid accumulating over time in the pseudocoelomic cavity of the animal. Particularly intriguing is the striking parallel this phenotype has to congenital hydrocephalus, a chronic condition caused by progressive build-up of cerebrospinal fluid in the brain cavities or ventricles. Importantly, hydrocephalus or its milder form, ventriculomegaly, is frequently observed in patients with L1 or Rasopathy syndrome caused respectively by genetic variants in *L1CAM* or in genes that result in elevated ERK signaling (Varagur *et al*. 2022; Weaver and GRIPP 2022; Aragon *et al*. 2024). Yet another point of interest is that mice homozygous for null mutations in the RanBPM-encoding *RanBP9* gene exhibit ventriculomegaly, similar to *L1cam* null mice (Dahme *et al*. 1997; Palavicini *et al*. 2013). Based on these studies, we predict that like in the developing vulva, GID-1 and SAX-7 likely function together in the *C. elegans* nervous system as well as in tissues regulating fluid homeostasis.

### SAX-7 as a general modulator of ERK activation

This study underscores the utility of genetic modifier studies to identify genetic functions that are less apparent because they are of a modulatory capacity or because of genetic compensation. In fact, it was in a genetic modifier screen that uncovered the genetically redundant role for *sax-7* in gastrulation (Grana *et al*. 2010). In our study, we used genetic modifier analyses to uncover a role for *sax-7* as a general modulator of MPK-1/ERK signaling in multiple tissues. This finding raises the question of whether this SAX-7 role is regulated; if so, what are the signals and are they shared in these disparate processes? Alternatively, could SAX-7 modulating ERK signaling in a constitutive manner, perhaps to maintain a consistently low level of activated MPK-1? A minimal baseline level of inactivated MPK-1 is critical to ensure sensitivity to small increases in activating signals and a quick cellular response to activated MPK-1. SAX-7 is localized to the plasma membrane of virtually all *C. elegans* cells, placing it in an ideal subcellular location to regulate conventional ERK signaling activated by receptor tyrosine kinases (Chen *et al*. 2001; Zhou *et al*. 2008; Moseley-ALldredge *et al*. 2022). Importantly, L1CAMs can biochemically interact with the FGF receptor, both directly and indirectly, in *C. elegans* and mammalian cells (Doherty and Walsh 1996; Saffell *et al*. 1997; Kulahin *et al*. 2008; DÍaz-Balzac *et al*. 2015). This is indeed intriguing considering the role of FGFR hyperactivity in fluid dysregulation and the Clr phenotype. Similarly, L1CAMs biochemically interact with EGF receptors (Donier *et al*. 2012; Chien *et al*. 2024), another point of interest relative to the functional intersection of SAX-7 and LET-23/EGFR in vulval development.

The ubiquitous expression of SAX-7, together with its numerous protein-protein motifs, leas us to speculate SAX-7 function as a scaffold to provide spatial specificity and dynamic regulation of diverse signaling complexes (Langeberg and Scott 2015); as such, SAX-7 likely has many as-yet-undiscovered roles. Indeed, SAX-7 has established interactions with diverse cytoplasmic, integral membrane, and extracellular proteins that allow SAX-7 to mediate the aforementioned neuronal functions as well as non-neuronal roles, including gastrulation, pharyngeal morphogenesis, and maintenance of epithelial apical junctions (Axang *et al*. 2007; Zhou *et al*. 2008; Grana *et al*. 2010; Zhou and Chen 2011; Lynch *et al*. 2012; Dong *et al*. 2013; Díaz-Balzac *et al*. 2015; Diaz-Balzac *et al*. 2016; Zou *et al*. 2016). As with the SAX-7 roles in modulating MPK-1/ERK signaling and in gastrulation, we suspect that genetic modifier studies will play a critical role in identifying additional SAX-7 functions, which may give insight into how L1CAMs contribute to associated polygenic disorders.

## ACKNOWLEDGEMENTS

We are grateful to the *C. elegans* Genetics Center, which is funded by the NIH office of Research Infrastructure Programs (P40 OD10440), for providing several strains used in the study. We thank the National BioResource Project Japan for providing the *gid-1(tm3703)* strain and David Greenstein and Melissa Gardner at the University of Minnesota for use of the Nikon Ti2 inverted confocal microscope and NIS elements (Nikon Inc., Melville, NY). This work was supported by NIH grant NS045873 to L.C.

## REFERENCES

Abdus-Saboor, I., V. P. Mancuso, J. I. Murray, K. Palozola, C. Norris et al., 2011 Notch and Ras promote sequential steps of excretory tube development in C. elegans. Development 138: 3545–3555.

Adams, J. P., A. E. Anderson, A. W. Varga, K. T. Dineley, R. G. Cook et al., 2000 The A-type potassium channel Kv4.2 is a substrate for the mitogen-activated protein kinase ERK. J Neurochem 75: 2277–2287.

Alfonso, A., K. Grundahl, J. R. Mcmanus and J. B. Rand, 1994 Cloning and characterization of the choline acetyltransferase structural gene (cha-1) from C. elegans. J Neurosci 14: 2290–2300.

Anderson, R. B., K. N. Turner, A. G. Nikonenko, J. Hemperly, M. Schachner et al., 2006 The cell adhesion molecule l1 is required for chain migration of neural crest cells in the developing mouse gut. Gastroenterology 130: 1221–1232.

Aragon, C., D. Robinson, M. Kocher, K. Barrick, L. Chen et al., 2024 Genetic etiologies and diagnostic methods for congenital ventriculomegaly and hydrocephalus: A scoping review. Birth Defects Res 116: e2287.

Aroian, R. V., and P. W. Sternberg, 1991 Multiple functions of let-23, a Caenorhabditis elegans receptor tyrosine kinase gene required for vulval induction. Genetics 128: 251–267.

Atabakhsh, E., and C. Schild-Poulter, 2012 RanBPM is an inhibitor of ERK signaling. PLoS One 7: e47803.

Axang, C., M. Rauthan, D. H. Hall and M. Pilon, 2007 The twisted pharynx phenotype in C. elegans. BMC Dev Biol 7: 61.

Ayalew, M., H. Le-Niculescu, D. F. Levey, N. Jain, B. Changala et al., 2012 Convergent functional genomics of schizophrenia: from comprehensive understanding to genetic risk prediction. Mol Psychiatry 17: 887–905.

Bastiani, C. A., S. Gharib, M. I. Simon and P. W. Sternberg, 2003 Caenorhabditis elegans Galphaq regulates egg-laying behavior via a PLCbeta-independent and serotonin-dependent signaling pathway and likely functions both in the nervous system and in muscle. Genetics 165: 1805–1822.

Beitel, G. J., S. G. Clark and H. R. Horvitz, 1990 Caenorhabditis elegans ras gene let-60 acts as a switch in the pathway of vulval induction. Nature 348: 503–509.

Berset, T., E. F. Hoier, G. Battu, S. Canevascini and A. Hajnal, 2001 Notch inhibition of RAS signaling through MAP kinase phosphatase LIP-1 during C. elegans vulval development. Science 291: 1055–1058.

Brennan, D. F., A. C. Dar, N. T. Hertz, W. C. Chao, A. L. Burlingame et al., 2011 A Raf-induced allosteric transition of KSR stimulates phosphorylation of MEK. Nature 472: 366–369.

Brenner, S., 1974 The genetics of Caenorhabditis elegans. Genetics 77: 71–94.

Cebul, E. R., I. G. Mclachlan and M. G. Heiman, 2020 Dendrites with specialized glial attachments develop by retrograde extension using SAX-7 and GRDN-1. Development 147.

Chen, L., Y. Fu, M. Ren, B. Xiao and C. S. Rubin, 2011 A RasGRP, C. elegans RGEF-1b, couples external stimuli to behavior by activating LET-60 (Ras) in sensory neurons. Neuron 70: 51–65.

Chen, L., B. Ong and V. Bennett, 2001 LAD-1, the Caenorhabditis elegans L1CAM homologue, participates in embryonic and gonadal morphogenesis and is a substrate for fibroblast growth factor receptor pathway-dependent phosphotyrosine-based signaling. J Cell Biol 154: 841–855.

Chen, L., and S. Zhou, 2010 “CRASH”ing with the worm: insights into L1CAM functions and mechanisms. Dev Dyn 239: 1490–1501.

Cheng, L., S. Lemmon and V. Lemmon, 2005 RanBPM is an L1-interacting protein that regulates L1-mediated mitogen-activated protein kinase activation. J Neurochem 94: 1102–1110.

Chien, M. H., Y. C. Yang, K. H. Ho, Y. F. Ding, L. H. Chen et al., 2024 Cyclic increase in the ADAMTS1-L1CAM-EGFR axis promotes the EMT and cervical lymph node metastasis of oral squamous cell carcinoma. Cell Death Dis 15: 82.

Coleman, B., I. Topalidou and M. Ailion, 2018 Modulation of Gq-Rho Signaling by the ERK MAPK Pathway Controls Locomotion in Caenorhabditis elegans. Genetics 209: 523–535.

Consortium, C. E. D. M., 2012 large-scale screening for targeted knockouts in the Caenorhabditis elegans genome. G3 (Bethesda) 2: 1415–1425.

Dahme, M., U. Bartsch, R. Martini, B. Anliker, M. Schachner et al., 1997 Disruption of the mouse L1 gene leads to malformations of the nervous system. Nat Genet 17: 346–349.

Das, S., S. Haq and S. Ramakrishna, 2018 Scaffolding protein RanBPM and its interactions in diverse signaling pathways in health and disease. Discov Med 25: 177–194.

de la Cova, C., R. Townley, S. Regot and I. Greenwald, 2017 A Real-Time Biosensor for ERK Activity Reveals Signaling Dynamics during C. elegans Cell Fate Specification. Dev Cell 42: 542–553 e544.

Desse, V. E., C. R. Blanchette, M. Nadour, P. Perrat, L. Rivollet et al., 2021 Neuronal postdevelopmentally acting SAX-7S/L1CAM can function as cleaved fragments to maintain neuronal architecture in Caenorhabditis elegans. Genetics 218.

Díaz-Balzac, C. A., M. I. Lázaro-Peña, G. A. Ramos-Ortiz and H. E. Bülow, 2015 The Adhesion Molecule KAL-1/anosmin-1 Regulates Neurite Branching through a SAX-7/L1CAM-EGL-15/FGFR Receptor Complex. Cell Rep 11: 1377–1384.

Diaz-Balzac, C. A., M. Rahman, M. I. Lazaro-Pena, L. A. Martin Hernandez, Y. Salzberg et al., 2016 Muscle- and Skin-Derived Cues Jointly Orchestrate Patterning of Somatosensory Dendrites. Curr Biol 26: 2379–2387.

Doherty, P., and F. S. Walsh, 1996 CAM-FGF receptor interactions: a model for axonal growth. Mol Cell Neurosci 8: 99–111.

Dong, X., O. W. Liu, A. S. Howell and K. Shen, 2013 An extracellular adhesion molecule complex patterns dendritic branching and morphogenesis. Cell 155: 296–307.

Donier, E., J. A. Gomez-Sanchez, C. Grijota-Martinez, J. Lakoma, S. Baars et al., 2012 L1CAM binds ErbB receptors through Ig-like domains coupling cell adhesion and neuregulin signalling. PLoS One 7: e40674.

Douglas, D. S., J. L. Moran, J. R. Bermingham, Jr., X. J. Chen, D. N. Brindley et al., 2009 Concurrent Lpin1 and Nrcam mouse mutations result in severe peripheral neuropathy with transitory hindlimb paralysis. J Neurosci 29: 12089–12100.

Duncan, B. W., K. E. Murphy and P. F. Maness, 2021 Molecular Mechanisms of L1 and NCAM Adhesion Molecules in Synaptic Pruning, Plasticity, and Stabilization. Front Cell Dev Biol 9: 625340.

Elahi, Z., M. Soveyzi, S. Nafissi, Y. Nilipour, M. Goleyjani Moghadam et al., 2023 Bi-allelic loss of function variant in the NRCAM gene is associated with motor-predominant axonal polyneuropathy; the second report. Mol Genet Genomic Med 11: e2131.

Frodyma, D., B. Neilsen, D. Costanzo-Garvey, K. Fisher and R. Lewis, 2017 Coordinating ERK signaling via the molecular scaffold Kinase Suppressor of Ras. F1000Res 6: 1621.

Gauntner, T. D., M. Karumuri, M. A. Guzman, S. E. Starnes, S. Besmer et al., 2021 Hirschsprung Disease in an Infant with L1 syndrome: Report of a New Case and a novel L1CAM variant. Clin Case Rep 9: 1518–1523.

Gauthier, K., and C. E. Rocheleau, 2017 C. elegans Vulva Induction: An In Vivo Model to Study Epidermal Growth Factor Receptor Signaling and Trafficking. Methods Mol Biol 1652: 43–61.

Grana, T. M., E. A. Cox, A. M. Lynch and J. Hardin, 2010 SAX-7/L1CAM and HMR-1/cadherin function redundantly in blastomere compaction and non-muscle myosin accumulation during Caenorhabditis elegans gastrulation. Dev Biol 344: 731–744.

Gronska-Peski, M., M. Schachner and J. M. Hebert, 2020 L1cam curbs the differentiation of adult-born hippocampal neurons. Stem Cell Res 48: 101999.

Guo, M. L., B. Xue, D. Z. Jin, L. M. Mao and J. Q. Wang, 2012 Interactions and phosphorylation of postsynaptic density 93 (PSD-93) by extracellular signal-regulated kinase (ERK). Brain Res 1465: 18–25.

Hamakawa, M., T. Uozumi, N. Ueda, Y. Iino and T. Hirotsu, 2015 A role for Ras in inhibiting circular foraging behavior as revealed by a new method for time and cell-specific RNAi. BMC Biol 13: 6.

Hortsch, M., K. Nagaraj and R. Mualla, 2014 The L1 family of cell adhesion molecules: a sickening number of mutations and protein functions. Adv Neurobiol 8: 195–229.

Huang, P., and M. J. Stern, 2004 FGF signaling functions in the hypodermis to regulate fluid balance in C. elegans. Development 131: 2595–2604.

Jacobs, D., G. J. Beitel, S. G. Clark, H. R. Horvitz and K. Kornfeld, 1998 Gain-of-function mutations in the Caenorhabditis elegans lin-1 ETS gene identify a C-terminal regulatory domain phosphorylated by ERK MAP kinase. Genetics 149: 1809–1822.

Jouet, M., A. Rosenthal, G. Armstrong, J. Macfarlane, R. Stevenson et al., 1994 X-linked spastic paraplegia (SPG1), MASA syndrome and X-linked hydrocephalus result from mutations in the L1 gene. Nat Genet 7: 402–407.

Kidger, A. M., and S. M. Keyse, 2016 The regulation of oncogenic Ras/ERK signalling by dual-specificity mitogen activated protein kinase phosphatases (MKPs). Semin Cell Dev Biol 50: 125–132.

Kiel, C., D. Matallanas and W. Kolch, 2021 The Ins and Outs of RAS Effector Complexes. Biomolecules 11.

Kim, W., R. S. Underwood, I. Greenwald and D. D. Shaye, 2018 OrthoList 2: A New Comparative Genomic Analysis of Human and Caenorhabditis elegans Genes. Genetics 210: 445–461.

Kortum, R. L., and R. E. Lewis, 2004 The molecular scaffold KSR1 regulates the proliferative and oncogenic potential of cells. Mol Cell Biol 24: 4407–4416.

Kulahin, N., S. Li, A. Hinsby, V. Kiselyov, V. Berezin et al., 2008 Fibronectin type III (FN3) modules of the neuronal cell adhesion molecule L1 interact directly with the fibroblast growth factor (FGF) receptor. Mol Cell Neurosci 37: 528–536.

Kurolap, A., F. Kreuder, C. Gonzaga-Jauregui, M. P. Duvdevani, T. Harel et al., 2022 Bi-allelic variants in neuronal cell adhesion molecule cause a neurodevelopmental disorder characterized by developmental delay, hypotonia, neuropathy/spasticity. Am J Hum Genet 109: 518–532.

Lackner, M. R., and S. K. Kim, 1998 Genetic analysis of the Caenorhabditis elegans MAP kinase gene mpk-1. Genetics 150: 103–117.

Lackner, M. R., K. Kornfeld, L. M. Miller, H. R. Horvitz and S. K. Kim, 1994 A MAP kinase homolog, mpk-1, is involved in ras-mediated induction of vulval cell fates in Caenorhabditis elegans. Genes Dev 8: 160–173.

Lackner, M. R., S. J. Nurrish and J. M. Kaplan, 1999 Facilitation of synaptic transmission by EGL-30 Gqalpha and EGL-8 PLCbeta: DAG binding to UNC-13 is required to stimulate acetylcholine release. Neuron 24: 335–346.

Lakshmana, M. K., C. D. Hayes, S. P. Bennett, E. Bianchi, K. M. Reddy et al., 2012 Role of RanBP9 on amyloidogenic processing of APP and synaptic protein levels in the mouse brain. FASEB J 26: 2072–2083.

Langeberg, L. K., and J. D. Scott, 2015 Signalling scaffolds and local organization of cellular behaviour. Nat Rev Mol Cell Biol 16: 232–244.

Levchenko, A., J. Bruck and P. W. Sternberg, 2000 Scaffold proteins may biphasically affect the levels of mitogen-activated protein kinase signaling and reduce its threshold properties. Proc Natl Acad Sci U S A 97: 5818–5823.

Lynch, A. M., T. Grana, E. Cox-Paulson, A. Couthier, M. Cameron et al., 2012 A genome-wide functional screen shows MAGI-1 is an L1CAM-dependent stabilizer of apical junctions in C. elegans. Curr Biol 22: 1891–1899.

Madeira, F., N. Madhusoodanan, J. Lee, A. Eusebi, A. Niewielska et al., 2024 The EMBL-EBI Job Dispatcher sequence analysis tools framework in 2024. Nucleic Acids Res 52: W521–W525.

Mao, L. M., and J. Q. Wang, 2016 Synaptically Localized Mitogen-Activated Protein Kinases: Local Substrates and Regulation. Mol Neurobiol 53: 6309–6315.

Martin, S. W., A. J. Butcher, N. S. Berrow, M. W. Richards, R. E. Paddon et al., 2006 Phosphorylation sites on calcium channel alpha1 and beta subunits regulate ERK-dependent modulation of neuronal N-type calcium channels. Cell Calcium 39: 275–292.

Mcmullan, R., E. Hiley, P. Morrison and S. J. Nurrish, 2006 Rho is a presynaptic activator of neurotransmitter release at pre-existing synapses in C. elegans. Genes Dev 20: 65–76.

Mctavish, C. J., W. Berube-Janzen, X. Wang, M. E. R. Maitland, L. M. Salemi et al., 2019 Regulation of c-Raf Stability through the CTLH Complex. Int J Mol Sci 20.

Miller, K. G., A. Alfonso, M. Nguyen, J. A. Crowell, C. D. Johnson et al., 1996a A genetic selection for Caenorhabditis elegans synaptic transmission mutants. Proc Natl Acad Sci U S A 93: 12593–12598.

Miller, K. G., M. D. Emerson and J. B. Rand, 1999 Goalpha and diacylglycerol kinase negatively regulate the Gqalpha pathway in C. elegans. Neuron 24: 323–333.

Miller, L. M., D. A. Waring and S. K. Kim, 1996b Mosaic analysis using a ncl-1 (+) extrachromosomal array reveals that lin-31 acts in the Pn.p cells during Caenorhabditis elegans vulval development. Genetics 143: 1181–1191.

Miningou, N., and K. T. Blackwell, 2020 The road to ERK activation: Do neurons take alternate routes? Cell Signal 68: 109541.

Monfrini, E., L. Straniero, S. Bonato, G. Monzio Compagnoni, A. Bordoni et al., 2019 Neurofascin (NFASC) gene mutation causes autosomal recessive ataxia with demyelinating neuropathy. Parkinsonism Relat Disord 63: 66–72.

Morelli, K. H., K. L. Seburn, D. G. Schroeder, E. L. Spaulding, L. A. Dionne et al., 2017 Severity of Demyelinating and Axonal Neuropathy Mouse Models Is Modified by Genes Affecting Structure and Function of Peripheral Nodes. Cell Rep 18: 3178–3191.

Moseley-Alldredge, M., S. Sheoran, H. Yoo, C. O’keefe, J. E. Richmond et al., 2022 A role for the Erk MAPK pathway in modulating SAX-7/L1CAM-dependent locomotion in Caenorhabditis elegans. Genetics 220.

Nguyen, A., W. R. Burack, J. L. Stock, R. Kortum, O. V. Chaika et al., 2002 Kinase suppressor of Ras (KSR) is a scaffold which facilitates mitogen-activated protein kinase activation in vivo. Mol Cell Biol 22: 3035–3045.

Ojea Ramos, S., M. Feld and M. S. Fustinana, 2022 Contributions of extracellular-signal regulated kinase 1/2 activity to the memory trace. Front Mol Neurosci 15: 988790.

Opperman, K., M. Moseley-Alldredge, J. Yochem, L. Bell, T. Kanayinkal et al., 2015 A novel nondevelopmental role of the sax-7/L1CAM cell adhesion molecule in synaptic regulation in Caenorhabditis elegans. Genetics 199: 497–509.

Palavicini, J. P., B. N. Lloyd, C. D. Hayes, E. Bianchi, D. E. Kang et al., 2013 RanBP9 Plays a Critical Role in Neonatal Brain Development in Mice. PLoS One 8: e66908.

Palavicini, J. P., H. Wang, D. Minond, E. Bianchi, S. Xu et al., 2014 RanBP9 overexpression down-regulates phospho-cofilin, causes early synaptic deficits and impaired learning, and accelerates accumulation of amyloid plaques in the mouse brain. J Alzheimers Dis 39: 727–740.

Parisi, M. A., R. P. Kapur, I. Neilson, R. M. Hofstra, L. W. Holloway et al., 2002 Hydrocephalus and intestinal aganglionosis: is L1CAM a modifier gene in Hirschsprung disease? Am J Med Genet 108: 51–56.

Robinson-Thiewes, S., B. Dufour, P. O. Martel, X. Lechasseur, A. A. D. Brou et al., 2021 Non-autonomous regulation of germline stem cell proliferation by somatic MPK-1/MAPK activity in C. elegans. Cell Rep 35: 109162.

Rodriguez Torres, C. S., N. B. Wicker, V. Puccini de Castro, M. Stefinko, D. C. Bennett, et al., 2024 The Caenorhabditis elegans protein SOC-3 permits an alternative mode of signal transduction by the EGL-15 FGF receptor. Dev Biol 516: 183–195.

Roskoski, R., Jr., 2012 ERK1/2 MAP kinases: structure, function, and regulation. Pharmacol Res 66: 105–143.

Roy, F., G. Laberge, M. Douziech, D. Ferland-Mccollough and M. Therrien, 2002 KSR is a scaffold required for activation of the ERK/MAPK module. Genes Dev 16: 427–438.

Saffell, J. L., E. J. Williams, I. J. Mason, F. S. Walsh and P. Doherty, 1997 Expression of a dominant negative FGF receptor inhibits axonal growth and FGF receptor phosphorylation stimulated by CAMs. Neuron 18: 231–242.

Sakurai, K., O. Migita, M. Toru and T. Arinami, 2002 An association between a missense polymorphism in the close homologue of L1 (CHL1, CALL) gene and schizophrenia. Mol Psychiatry 7: 412–415.

Sakurai, T., 2012 The role of NrCAM in neural development and disorders--beyond a simple glue in the brain. Mol Cell Neurosci 49: 351–363.

Salemi, L. M., M. E. R. Maitland, C. J. Mctavish and C. Schild-Poulter, 2017 Cell signalling pathway regulation by RanBPM: molecular insights and disease implications. Open Biol 7.

Schrader, L. A., S. G. Birnbaum, B. M. Nadin, Y. Ren, D. Bui et al., 2006 ERK/MAPK regulates the Kv4.2 potassium channel by direct phosphorylation of the pore-forming subunit. Am J Physiol Cell Physiol 290: C852–861.

Shaltout, T. E., K. A. Alali, S. Bushra, A. M. Alkaseri, E. D. Jose et al., 2013 Significant association of close homologue of L1 gene polymorphism rs2272522 with schizophrenia in Qatar. Asia Pac Psychiatry 5: 17–23.

Shaye, D. D., and I. Greenwald, 2011 OrthoList: a compendium of C. elegans genes with human orthologs. PLoS One 6: e20085.

Shin, H., and D. J. Reiner, 2018 The Signaling Network Controlling C. elegans Vulval Cell Fate Patterning. J Dev Biol 6.

Smigiel, R., D. L. Sherman, M. Rydzanicz, A. Walczak, D. Mikolajkow et al., 2018 Homozygous mutation in the Neurofascin gene affecting the glial isoform of Neurofascin causes severe neurodevelopment disorder with hypotonia, amimia and areflexia. Hum Mol Genet 27: 3669–3674.

Smith, M. J., 2023 Defining bone fide effectors of RAS GTPases. Bioessays 45: e2300088.

Sternberg, P. W., 2005 Vulval development. WormBook: 1–28.

Sternberg, P. W., A. Golden and M. Han, 1993 Role of a raf proto-oncogene during Caenorhabditis elegans vulval development. Philos Trans R Soc Lond B Biol Sci 340: 259–265.

Sternberg, P. W., K. Van Auken, Q. Wang, A. Wright, K. Yook et al., 2024 WormBase 2024: status and transitioning to Alliance infrastructure. Genetics 227.

Sundaram, M. V., 2013 Canonical RTK-Ras-ERK signaling and related alternative pathways. WormBook: 1–38.

Sundaram, M. V., and M. Buechner, 2016 The Caenorhabditis elegans Excretory System: A Model for Tubulogenesis, Cell Fate Specification, and Plasticity. Genetics 203: 35–63.

Sundararajan, L., J. Stern and D. M. Miller, 3rd, 2019 Mechanisms that regulate morphogenesis of a highly branched neuron in C. elegans. Dev Biol 451: 53–67.

Sweatt, J. D., 2004 Mitogen-activated protein kinases in synaptic plasticity and memory. Curr Opin Neurobiol 14: 311–317.

Tam, G. W., L. N. Van De Lagemaat, R. Redon, K. E. Strathdee, M. D. Croning et al., 2010 Confirmed rare copy number variants implicate novel genes in schizophrenia. Biochem Soc Trans 38: 445–451.

Tan, P. B., M. R. Lackner and S. K. Kim, 1998 MAP kinase signaling specificity mediated by the LIN-1 Ets/LIN-31 WH transcription factor complex during C. elegans vulval induction. Cell 93: 569–580.

Taneera, J., S. Lang, A. Sharma, J. Fadista, Y. Zhou et al., 2012 A systems genetics approach identifies genes and pathways for type 2 diabetes in human islets. Cell Metab 16: 122–134.

Thomas, G. M., and R. L. Huganir, 2004 MAPK cascade signalling and synaptic plasticity. Nat Rev Neurosci 5: 173–183.

Tomida, T., S. Oda, M. Takekawa, Y. Iino and H. Saito, 2012 The temporal pattern of stimulation determines the extent and duration of MAPK activation in a Caenorhabditis elegans sensory neuron. Sci Signal 5: ra76.

Turner, K. N., M. Schachner and R. B. Anderson, 2009 Cell adhesion molecule L1 affects the rate of differentiation of enteric neurons in the developing gut. Dev Dyn 238: 708–715.

Van Camp, G., E. Fransen, L. Vits, G. Raes and P. J. Willems, 1996 A locus-specific mutation database for the neural cell adhesion molecule L1CAM (Xq28). Hum Mutat 8: 391.

Varagur, K., S. A. Sanka and J. M. Strahle, 2022 Syndromic Hydrocephalus. Neurosurg Clin N Am 33: 67–79.

Vits, L., G. Van Camp, P. Coucke, E. Fransen, K. De Boulle et al., 1994 MASA syndrome is due to mutations in the neural cell adhesion gene L1CAM. Nat Genet 7: 408–413.

Vos, Y. J., and R. M. Hofstra, 2010 An updated and upgraded L1CAM mutation database. Hum Mutat 31: E1102–1109.

Wallace, A. S., C. Schmidt, M. Schachner, M. Wegner and R. B. Anderson, 2010 L1cam acts as a modifier gene during enteric nervous system development. Neurobiol Dis 40: 622–633.

Wang, X., J. Kweon, S. Larson and L. Chen, 2005 A role for the C. elegans L1CAM homologue lad-1/sax-7 in maintaining tissue attachment. Dev Biol 284: 273–291.

Wang, X. J., and M. Camilleri, 2019 Hirschsprung disease: Insights on genes, penetrance, and prenatal diagnosis. Neurogastroenterol Motil 31: e13732.

Weaver, K. N., and K. W. Gripp, 2022 Central nervous system involvement in individuals with RASopathies. Am J Med Genet C Semin Med Genet 190: 494–500.

Wu, Y., and D. J. Reiner, 2022 A signalling cascade for Ral. Small GTPases 13: 128–135.

Yochem, J., M. Sundaram and M. Han, 1997 Ras is required for a limited number of cell fates and not for general proliferation in Caenorhabditis elegans. Mol Cell Biol 17: 2716–2722.

Zand, T. P., D. J. Reiner and C. J. Der, 2011 Ras effector switching promotes divergent cell fates in C. elegans vulval patterning. Dev Cell 20: 84–96.

Zhong, X., J. Drgonova, C. Y. Li and G. R. Uhl, 2015 Human cell adhesion molecules: annotated functional subtypes and overrepresentation of addiction-associated genes. Ann N Y Acad Sci 1349: 83–95.

Zhou, S., and L. Chen, 2011 Neural integrity is maintained by dystrophin in C. elegans. J Cell Biol 192: 349–363.

Zhou, S., K. Opperman, X. Wang and L. Chen, 2008 unc-44 Ankyrin and stn-2 gamma-syntrophin regulate sax-7 L1CAM function in maintaining neuronal positioning in Caenorhabditis elegans. Genetics 180: 1429–1443.

Zou, W., A. Shen, X. Dong, M. Tugizova, Y. K. Xiang et al., 2016 A multi-protein receptor-ligand complex underlies combinatorial dendrite guidance choices in C. elegans. Elife 5.

